# MTMR regulates KRAS function by controlling plasma membrane levels of phospholipids

**DOI:** 10.1101/2024.01.22.576612

**Authors:** Taylor E. Lange, Ali Naji, Ransome van der Hoeven, Hong Liang, Yong Zhou, Gerald R.V. Hammond, John F. Hancock, Kwang-jin Cho

## Abstract

KRAS, a small GTPase involved in cell proliferation and differentiation, frequently gains activating mutations in human cancers. For KRAS to function, it must bind the plasma membrane (PM) via interactions between its membrane anchor and phosphatidylserine (PtdSer). Therefore, depleting PM PtdSer abrogates KRAS PM binding and activity. From a genome-wide siRNA screen to identify genes regulating KRAS PM localization, we identified a set of phosphatidylinositol (PI) 3-phosphatases: myotubularin-related proteins (MTMR) 2, 3, 4, and 7. Here, we show that silencing *MTMR 2/3/4/7* disrupts KRAS PM interactions by reducing PM PI 4-phosphate (PI4P) levels, thereby disrupting the localization and operation of ORP5, a lipid transfer protein maintaining PM PtdSer enrichment. Concomitantly, silencing *MTMR 2/3/4/7* elevates PM PI3P levels while reducing PM and total PtdSer levels. We also observed MTMR 2/3/4/7 expression is interdependent. We propose that the PI 3-phosphatase activity of MTMR is required for generating PM PI, necessary for PM PI4P synthesis, promoting the PM localization of PtdSer and KRAS.

**eTOC summary:** We discovered that silencing the phosphatidylinositol (PI) 3-phosphatase, *MTMR*, disrupts the PM localization of PtdSer and KRAS. We propose a model, where *MTMR* loss depletes PM PI needed for PM PI4P synthesis, an essential phospholipid for PM PtdSer enrichment, thereby impairing KRAS PM localization.

## Introduction

H-, N- and KRAS proteins are small GTPases that oscillate between inactive GDP-bound and active GTP-bound conformational states and operate in signaling cascades that control cell growth and proliferation (Gorfe and Cho, 2019; Henkels et al., 2021). Consistent with this key regulatory role, activating mutations of RAS are present in ∼20% of human cancers, with the majority occurring in the KRAS isoform (Prior et al., 2020). All RAS isoforms must be localized to the inner leaflet of the plasma membrane (PM) and be spatially organized into nanoclusters for biological activity (Gorfe and Cho, 2019; Henkels et al., 2021). Membrane interactions are mediated by the RAS C-terminal membrane anchor (Zhou et al., 2017). In the case of KRAS4B, the major expressed splice variant of KRAS (Hood et al., 2023) (hereafter KRAS), this anchor comprises a C-terminal farnesyl cysteine methyl ester, generated by posttranslational modification, and an adjacent polybasic domain of six continuous lysines (Gutierrez et al., 1989; Hancock et al., 1991; Hancock et al., 1989; Hancock et al., 1990). Recent work has shown that the KRAS membrane anchor encodes exquisite binding specificity for phosphatidylserine (PtdSer) lipids with one saturated and one desaturated acyl chain (Zhou et al., 2017). This lipid binding specificity is effectively hard wired into the anchor structure and renders KRAS PM binding, nanoclustering and hence biological function absolutely dependent on PM PtdSer content (Cho et al., 2012b; Cho et al., 2013; van der Hoeven et al., 2013; van der Hoeven et al., 2018).

PtdSer is synthesized in the ER and delivered to the PM against a concentration gradient by the lipid transfer proteins (LTPs), oxysterol-binding protein-related protein (ORP) 5 and 8, operating at ER-PM membrane contact sites. One molecule of phosphatidylinositol (PI) 4-phosphate (P), generated by PM localized PI 4-kinase IIIα (PI4KA) from PI, is exchanged by ORP5/8 for one molecule of ER localized PtdSer (Chung et al., 2015; Moser von Filseck et al., 2015). PI4P delivered to the ER is converted to PI by the SAC1 phosphatase, which operating together with PI4KA, maintains the PI4P gradient that in turn, drives PtdSer transport to the PM (Chung et al., 2015; Moser von Filseck et al., 2015). Concordantly genetic and pharmacological studies have shown that inhibition or loss of any component of the PtdSer transport machinery, including PI4KA, EFR3A (the PM anchor protein for the kinase), ORP5, ORP8 and SAC1 results in a reduction in PM PtdSer content, displacement of KRAS from the PM and abrogation of KRAS signaling (Gulyas et al., 2017; Kattan et al., 2019; Kattan et al., 2021; Sohn et al., 2018). Other pharmacological approaches to deplete PM PtdSer also dissociate KRAS from the PM (Cho et al., 2012b; Garrido et al., 2020; Miller et al., 2019; Tan et al., 2019; Tan et al., 2018; van der Hoeven et al., 2018). The PM is a dynamic organelle, thus KRAS on endomembranes is sequestered by a chaperone protein, phosphodiesterase 6δ, and released to perinuclear membranes in the ARL2/3-dependent manner. The negative charge on the recycling endosome membranes then electrostatically traps KRAS, from where vesicular transport returns KRAS to the PM (Chandra et al., 2012; Schmick et al., 2014). KRAS phosphorylation at the Ser181 residue can disrupt this electrostatic interaction, depleting KRAS from the PM (Cho et al., 2016a; Kovar et al., 2020).

In this context we carried out a cell-based high content screen using a human genomic short interfering (siRNA) library to identify additional potential regulators of KRAS PM localization. Among the hits from the assay were multiple genes that encode proteins with phosphatase activity. Four of these genes encode myotubularin-related proteins (MTMR) 2, 3, 4 and 7, which have 3-phosphatase activity towards phosphatidylinositol (PI) 3-phosphate (P) and PI(3,5)-bisphosphate (PI(3,5)P_2_) (Robinson and Dixon, 2006). In this study, we examined the role of MTMR in the PM localization of KRAS. Our data demonstrate that knockdown (KD) of MTMR 2, 3, 4 or 7 dissociates KRAS, but not HRAS from the PM. We show *inter alia* that the molecular mechanism involves depletion of the PI4P and PtdSer content of the PM, extending the repertoire of proteins required to support the operation of the ORP5/8 LTPs and hence sustain KRAS PM localization.

## Method and Material

### Plasmids and reagents

shRNA glycerol sets for MTMR2 (RHS4533-EG8898), MTMR3 (RHS4533-EG8897), MTMR4 (RHS4533-EG9110), MTMR7 (RHS4533-EG9108), and human cDNA of MTMR2 (MHS6278-202760229), MTMR3 (MHS6278-202800229), MTMR4 (MHS6278-202801286) and MTMR7 (MHS6278-211688199) were purchased from GE Healthcare Dharmacon. Antibodies were purchased for detecting MTMR2 (PA5-22748; 1:1,000) and c-MYC (13-2500, 1:1,000) from Invitrogen, MTMR3 (sc-393779; 1:1,000) from SCBT, and MTMR7 (AB150458; 1:1,000) from Abcam. Antibodies for detecting c-MYC (10828-1-AP, 1:3,000), β-actin (60008-1-Ig; 1:5,000), MTMR3 (21336-1-AP; 1:1,000) and GFP (66002-1-lg; 1:4,000) were purchased from Proteintech. Anti-ppERK (4370; 1:3,000), anti-pAkt (S473) (4060; 1:1,000), anti-total ERK (4696; 1:1,000), anti-total Akt (2920; 1:1,000) antibodies for immunoblotting were from Cell Signaling Technology. Anti-RFP antibody (sc-390909, 1:1,000) was from Santa Cruz. Goat anti-mouse IgG (G21040; 1:2,000), anti-rabbit IgG secondary antibodies (G21234; 1:5,000) and CellMask Deep Red plasma membrane stain (C10046; 1:5,000) were purchased from Invitrogen. Staurosporine (BIA-S1086) was purchased from BioAustralis.

### Human whole-genome siRNA screen

A whole-genome siRNA screen was performed in Caco-2 cells stably expressing GFP-KRASG12V using a pre-aliquoted Silencer siRNA library (Ambion) at 30 nM final concentration. A total of 21,687 genes were represented in the library and were arrayed into glass-bottomed sixty-nine 384-well plates in triplicate. Gene silencing was induced for 4 days via reverse transfection using Dharmafect 1 (GE Healthcare). For controls, each plate included non-targeting control siRNA and siRNA targeting *FNTA* (farnesyltransferase type1 subunit alpha) as negative and positive controls, respectively. Also, each plate included the *Kif11* siRNA that targets a gene essential for cell survival served as control for transfection and knockdown efficiency. After the gene silencing, cells were fixed with 4% paraformaldehyde (PFA) and stored in 1x phosphate-buffered saline (PBS) at 4°C until imaged. Plates were imaged using Nikon A1R confocal microscope. Images were acquired as a 2 by 1 montage with a 20x objective lens using the GFP-confocal mode with NIS-Elements automated imaging software. Images were scored for KRAS PM dissociation/intracellular accumulation of GFP- KRASG12V and cytotoxicity. 253 genes were selected from the primary screen based on KRAS PM dissociation, and after a counter screen with GFP-HRASG12V to identify KRAS-specific mediators, we refined the candidate list to 157. The 157 genes were further validated with an independent secondary screen involving silencing each gene with Dharmacon SmartPool ON-TARGET plus siRNA in Caco-2 cells expressing GFP-KRASG12V and scoring for KRAS PM dissociation. A further bioinformatics analysis revealed that 8 of these genes encode proteins with phosphatase activity (Table 1).

**Table 1.** 8 novel genes, which encode proteins with phosphatase activity are identified to regulate KRAS/PM localization. PI – phosphatidylinositol, Ins – inositol, MTMR – myotubularin-related protein, MINPP1 – multiple inositol-polyphosphate phosphatase 1, TMEM55A – transmembrane protein 55A, INPPL1 - inositol polyphosphate phosphatase Like 1, SHIP2 – SH2-containing phosphatidylinositol 3,4,5-trisphosphate 5-phosphatase, PPP2R5B - protein phosphatase 2 regulatory subunit β

| Protein Name | Localization | Target PIs | Ref. |
| --- | --- | --- | --- |
| MTMR2 | Perinuclear, cytoplasm | PI 3-phosphatase for PI3P and PI(3,5)P<sub>2</sub> | (Kim et al., 2003), (Cao et al., 2008) |
| MTMR3 | ER, cytoplasm | PI 3-phosphatase for PI3P and PI(3,5)P<sub>2</sub> | (Walker et al., 2001), (Lorenzo et al., 2005) |
| MTMR4 | Early and recycling endosomes | PI 3-phosphatase for PI3P | (Naughtin et al., 2010), (Pham et al., 2018) |
| MTMR7 | Golgi complex, endosomes, cytoplasm | PI 3-phosphatase for PI3P | (Mochizuki and Majerus, 2003) |
| MINPP1 | ER | 3-phosphatase for Ins(1,3,4,5,6)pentakisphosphate (InsP<sub>5</sub>), inositol hexakisphosphate (InsP<sub>6</sub>) and 2,3bisphosphoglycerate | (Chi et al., 2000), (Cho et al., 2008) |
| TMEM55A | Lysosome, late endosome | PI 4-phosphatase for PI(4,5)P<sub>2</sub> | (Ungewickell et al., 2005) |
| INPPL1/SHIP2 | Cytoplasm, plasma membrane, cytoskeleton, lamellipodia | PI 5-phosphatase for PI(3,4,5)P<sub>3</sub> | (Taylor et al., 2000), (Dyson et al., 2001) |
| PPP2R5B | Cytoplasm | A component of the serine/threonine-protein phosphatase 2A complex (PP2A). Mediates PP2A dephosphorylation of Akt. | (McCright et al., 1996), (Rodgers et al., 2011) |

### Cell lines

T47D (HTB-13, ATCC) and BxPC3 (CRL-1687, ATCC) cells were maintained in RPMI-1640 medium (30-2001, ATCC) supplemented with 2 mM L-glutamine (CA009-010, GenDepot) and 10% fetal bovine serum (FBS) (16000044, Gibco). For T47D cells, 10 μg/mL human recombinant insulin (12585-014, Gibco) was also supplemented. MIA PaCa-2 (CRL- 1420, ATCC), PANC-1 (CRL-1469, ATCC) and Caco-2 (HTB-37, ATCC) cells were maintained in DMEM medium with high glucose (CM002, Gibco) supplemented with 2 mM L-glutamine and 10% FBS. All cells were grown at 37°C with 5% CO_2_, and frequently tested for mycoplasma using MycoAlert Mycoplasma Detection Kit (LT07-318, Lonza).

### Generating T47D stable cell lines

T47D cells were transfected with GFP-KRASG12V, - HRASG12V or -LactC2 in pEF6 vector (Invitrogen) using *Trans*IT-BrCa Transfection Reagent (MIR 5500, Mirus), and selected with 10 μg/mL blasticidin for 3 weeks. After the selection, cells were maintained in 5 μg/mL blasticidin. For generating MTMR knockdown cell lines, wild-type (WT) T47D cells were infected with lentivirus expressing shRNA targeting *MTMR 2/3/4/7.* Cells were selected by puromycin (1 μg/mL) for 1 week and maintained in 0.5 μg/mL puromycin.

### Preparing T47D cells for imaging and calculating Manders’ coefficient

5 x 10^5^ T47D cells were seeded on a coverslip on day 1 in RPMI-1640 medium containing 10% FBS, 2 mM L-glutamine and 10 μg/mL insulin (complete growth medium). On day 2, fresh complete growth medium was supplemented. On day 3, cells were fixed with 4% PFA for 30 min, quenched with 50 mM NH_4_Cl for 10 min and mounted with Vectashield (H-1000, Vector Laboratories). For calculating Manders’ coefficient, cells were incubated with CellMask stain (1:5,000) for 1 h in complete growth medium, fixed with 4% PFA, and imaged by confocal microscopy. Using ImageJ software 1.53k, images were converted to 8-bit, and a threshold to a control pixel of each image was set. The fraction of red-fluorophore-conjugated CellMask co-localizing with GFP-tagged proteins was calculated using Manders’ coefficient plugin downloaded from Wright Cell Image Facility.

### Lipidomic analysis

T47D cells stably expressing GFP-KRASG12V with *MTMR 2/3/4/7* knockdown were cultured in complete growth medium containing 0.5 µg/ml puromycin. Triplicate samples of 1x10^6^ cells were prepared in 333 μL Dulbecco’s PBS (DPBS without Ca^2+^ and Mg^2+^, 14190144, Invitrogen). Lipid extraction and analysis using electron spray ionization and MS/ MS were performed at Lipotype GmbH (Dresden, Germany), as described previously (Gerl et al., 2012; Sampaio et al., 2011). Automated processing of acquired mass spectra, identification, and quantification of detected lipid species were done by LipidXplorer software. Only lipid identifications with a signal-to-noise ratio >5, an absolute abundance of at least 1 pmol, and a signal intensity 5-fold higher than in corresponding blank samples were considered for further data analysis. The abundance of lipids is represented as a heat map with log_2_ scale relative to control (scrambled shRNA) cells.

### Measuring total PI3P level

The total amount of PI(3)P was examined in a quantitative and competitive ELISA format assay according to the manufacturer’s instructions (K-3300, Echelon Biosciences). Briefly, *MTMR 2/3/4/ KD* T47D cells grown on T75 flasks were collected in ice-cold 0.5 M trichloroacetic acid (TCA) and centrifuged at 1,000 x g for 10 min at 4°C. The pellet was washed with 3 mL 5% TCA/1 mM EDTA and neutral lipids were removed using MeOH:CHCl_3_ (2:1), followed by acidic lipids extraction using MeOH:CHCl_3_:12N HCl (80:40:1). The acidic lipid extracts were further separated into organic and aqueous phases, and the organic phase was dried in a vacuum dryer. The dried lipid was rehydrated in PBS-T 3% protein stabilizer and sonicated in a water bath. The sample was then used to measure PI3P level using the mass ELISA kit. Absorbance was read at 450 nm on Synergy H1 Plate Reader (BioTek).

### Western blotting

Preparation of cell lysates and immunoblotting were performed as described previously (Cho et al., 2011). Briefly, cells were washed twice with ice-cold 1x PBS. Cells were harvested in lysis buffer B containing 50 mM Tris-Cl (pH 7.5), 75 mM NaCl, 25 mM NaF, 5 mM MgCl_2_, 5 mM EGTA, 1 mM DTT, 100 μM NaVO_4_, 1% NP-40 plus protease and phosphatase inhibitors. SDS-PAGE and immunoblotting were generally performed using 20 – 40 μg of lysate from each sample group. Signals were detected by enhanced chemiluminescence (34578 and 34075, Thermo Fisher Scientific) and imaged using an Amersham Imager 600 (GE Healthcare). ImageJ software (v1.53k) was used to quantify band intensity.

### Validating MTMR4 knockdown

Messenger RNA (mRNA) was extracted from T47D cells infected with lentivirus expressing shRNA targeting MTMR4 using the RNeasy kit (74104, Qiagen) according to the manufacturer’s instructions. 2 μg of total RNA was converted to cDNA using Superscript II reverse transcriptase (18064-014, Thermo Fisher Scientific). To verify knockdown, forward and reverse primers specific for human *MTMR4* exons 1 and 5, and *GAPDH* (glyceraldehyde-3-phosphate dehydrogenase) exons 1 and 3 were designed: for *MTMR4*, 5’-CCAAGCCAAGGATCTGTTCCC-3’ and 5’-TGTGTGAGACTCTCCAGACGT-‘3’, and for *GAPDH*, 5’- GGAGCGAGATCCCTCCAAAAT-3’ and 5’-GGCTGTTGTCATACTTCTCATGG-3’, respectively. cDNA was amplified by PCR (P2311, GenDepot) and the products were resolved by 1.5% agarose gel electrophoresis and visualized by ethidium bromide staining.

### *C. elegans* vulva quantification assay

*let-60* (n1046) worms were kindly provided by Swathi Arur (MD Anderson Cancer Center, Houston, TX). RNAi was induced by feeding *let-60* worms through adult stage with *E. coli* HT115, producing double-stranded RNA to target genes. The presence of the multivulva phenotype was scored using a DIC/Nomarski microscope. All RNAi clones were from the *C. elegans* RNAi (Ahringer) collection (Source BioScience) and were sequenced.

### Proliferation assay

*MTMR 2/3/4/7* knockdown T47D cells stably expressing GFP-KRASG12V or -HRASG12V (1 x 10^4^ per well), or human pancreatic ductal adenocarcinoma cell lines, BxPC3, MIA PaCa-2 and PANC-1 (3 x 10^4^ per well) were seeded in triplicate onto a 96-well plate in complete growth medium containing 1.0 μg/mL puromycin. Cells were cultured for 4 days, and fresh complete growth medium containing 1.0 μg/mL puromycin was supplemented every 24 h. On day 5, cell proliferation was assayed by counting cells using Countess II cell counter (Invitrogen).

### Annexin V binding assay

*MTMR 2/3/4/7* knockdown T47D cells stably expressing GFP- KRASG12V were incubated with Cy5-conjugated annexin V (559934, BD Pharmingen) according to the manufacturer’s instructions. Annexin V-positive cells were counted using BD AccuriC6 Analyzer with a 670LP nm filter. GFP-KRASG12V T47D cells expressing scrambled shRNA were treated with 1 μM staurosporine for 6 h to induce apoptosis.

### Electron microscopy

Plasma membrane (PM) sheets were prepared from Caco-2 or T47D cells expressing GFP-tagged proteins of interest and fixed as previously described (Hancock and Prior, 2005; Prior et al., 2003a; Prior et al., 2003b). For univariate analysis, PM sheets were labeled with anti-GFP antibody conjugated to 4.5-nm gold particles. Digital images of the immunogold-labeled PM sheets were taken in a transmission electron microscope. Intact 1 μm^2^ areas of the PM sheet were identified using ImageJ software (v1.53k), and the (x, y) coordinates of the gold particles were determined (Hancock and Prior, 2005; Prior et al., 2003a; Prior et al., 2003b). K-functions (Ripley, 1977) were calculated and standardized on the 99% confidence interval (CI) for univariate functions (Diggle, 2000; Hancock and Prior, 2005; Prior et al., 2003a; Prior et al., 2003b). In the case of univariate functions, a value of *L(r) – r* greater than the CI indicates significant clustering, and the maximum value of the function (L_max_) estimates the extent of clustering. Differences between replicated point patterns were analyzed by constructing bootstrap tests as described previously (Diggle, 2000; Plowman and Hancock, 2005), and the statistical significance against the results obtained with 1,000 bootstrap samples was evaluated.

### Confocal microscope imaging acquisition

A) Make and Model of microscope: Olympus Fluoview F1000. B) Type, magnification and numerical aperture of the objective lenses: Olympus UPlanSApo 60x/1.35 oil, ∞/0.17/FN26.5 UIS2 BFP1. C) Temperature: room temperature. D) Imaging medium: Vectashield (H-1000, Vector Laboratories). E) Fluorochromes: GFP and mCherry. F) Camera make and model: N/A. G) Acquisition software: FV10-ASW ver.4.2c. H) Any software used for imaging processing subsequent to data acquisition: None.

### A summary of Supplement Figures

Supplement Figure 1. *MTMR* 2/3/4/7 regulate the PM localization of KRASG12V.

Supplement Figure 2. *MTMR* 2/3/4/7 KD disrupts the PM localization and nanoclustering of CTK.

Supplement Figure 3. *MTMR* 2/3/4/7 regulate cellular PI3P contents.

Supplement Figure 4. Ectopic expression of WT *MTMR4* restores the effects of *MTMR4* loss.

## Results

### Genome-wide siRNA screening identifies novel genes regulating KRAS PM interaction

To identify novel regulators of KRAS PM interaction, we performed an image-based high content screen of a human genomic siRNA library. Caco-2 (human colorectal adenocarcinoma) cell line stably expressing monomeric green fluorescent protein (GFP)-tagged oncogenic mutant KRAS (KRASG12V) was generated and transfected with siRNA pools comprising four different oligonucleotides against each gene. Four days after transfection, cells were imaged using an automated confocal microscope to analyze the extent of KRAS mislocalization from the PM. We then narrowed the list of hits using a counter screen against HRASG12V to identify KRASG12V- specific mediators. Bioinformatics analysis revealed that 8 of these putative KRASG12V-specific mediators encode proteins with phosphatase activity (Table 1), 4 of which are members of the myotubularin-related protein (MTMR) family. There are 16 human myotubularins, which exhibit 3-phosphatase activity towards PI3P and PI(3,5)P_2_, generating PI and PI5P, respectively (Robinson and Dixon, 2006; Wang et al., 2024). MTMR family has previously been shown to regulate many cellular processes including endocytosis, membrane trafficking, cell proliferation, autophagy, cytokinesis and cytoskeletal dynamics (Hnia et al., 2012). We selected this *MTMR* gene set for further analysis.

### MTMR 2/3/4/7 regulate the PM localization of KRASG12V, but not HRASG12V

To validate the siRNA screen, we generated short-hairpin RNA (shRNA)-mediated stable KD cell lines. T47D (human mammary gland ductal carcinoma) epithelial cells stably expressing GFP- KRASG12V were infected with lentiviruses expressing shRNA targeting *MTMR2, 3, 4* and *7*, followed by puromycin selection. For each target gene, we tested 4 different shRNAs and selected the two most effective KD for further experiments. Immunoblotting verified that endogenous expression levels of MTMR2, 3 and 7 were reduced by the cognate shRNAs (Fig. 1A). MTMR4 mRNA levels were also reduced by shRNA expression; we evaluated mRNA because the anti-MTMR4 antibodies we tested were not able to detect endogenous MTMR4 (Fig. 1A). To study KRAS cellular localization, cells were incubated with CellMask, a dye staining cellular membranes (Hu et al., 2013; Maechler et al., 2019; Monkemoller et al., 2015; Park et al., 2019), and imaged by confocal microscopy. To quantify the extent of KRAS distribution to endomembranes, we calculated Manders’ coefficient, which measures the fraction of CellMask co-localized with GFP-KRASG12V (Cho et al., 2012b; Manders et al., 1993; Rehl et al., 2023). Our data show that while GFP-KRASG12V was predominantly localized to the PM in control cells, it was distributed intracellularly in the KD cell lines (Fig. 1B and S1A). Manders’ coefficients for CellMask are correspondingly higher in the KD cell lines, indicating that KRASG12V is redistributed to endomembranes after silencing *MTMR 2/3/4/7*. To directly quantify the extent of KRAS dissociation from the PM, intact basal PM sheets from T47D stably co-expressing GFP-KRASG12V and shRNA for *MTMR 2/3/4/7* were prepared and labeled with gold-conjugated anti-GFP antibodies and analyzed by electron microscopy (EM). We observed significant reductions in anti-GFP immunogold labeling after the KD, indicating loss of KRASG12V from the inner PM leaflet (Fig. 1C and S1B). Spatial mapping of GFP-KRASG12V on the PM in *MTMR 2/3/4/7* KD cells revealed significant decreases in *L_max_*, the peak value of the *L(r)-r* clustering statistic that measures the extent of KRAS nanoclustering, which is essential for high-fidelity RAS signal transduction (Cho and Hancock, 2013; Cho et al., 2012a; Tian et al., 2007) (Fig. 1E). Confocal and electron microscopy further demonstrated that while *MTMR 2/3/4/7* KD did not disrupt the PM localization and nanoclustering of HRASG12V, it disrupts that of the C-terminal truncated mutant KRAS (CTK), which contains the membrane anchor without the G-domain, from the PM (Fig. 1B, D, F and S2). Together, our data show that MTMR 2/3/4/7 regulate the PM localization and nanoclustering of KRAS, but not HRAS, independent of the G-domain.

**Figure 1.**
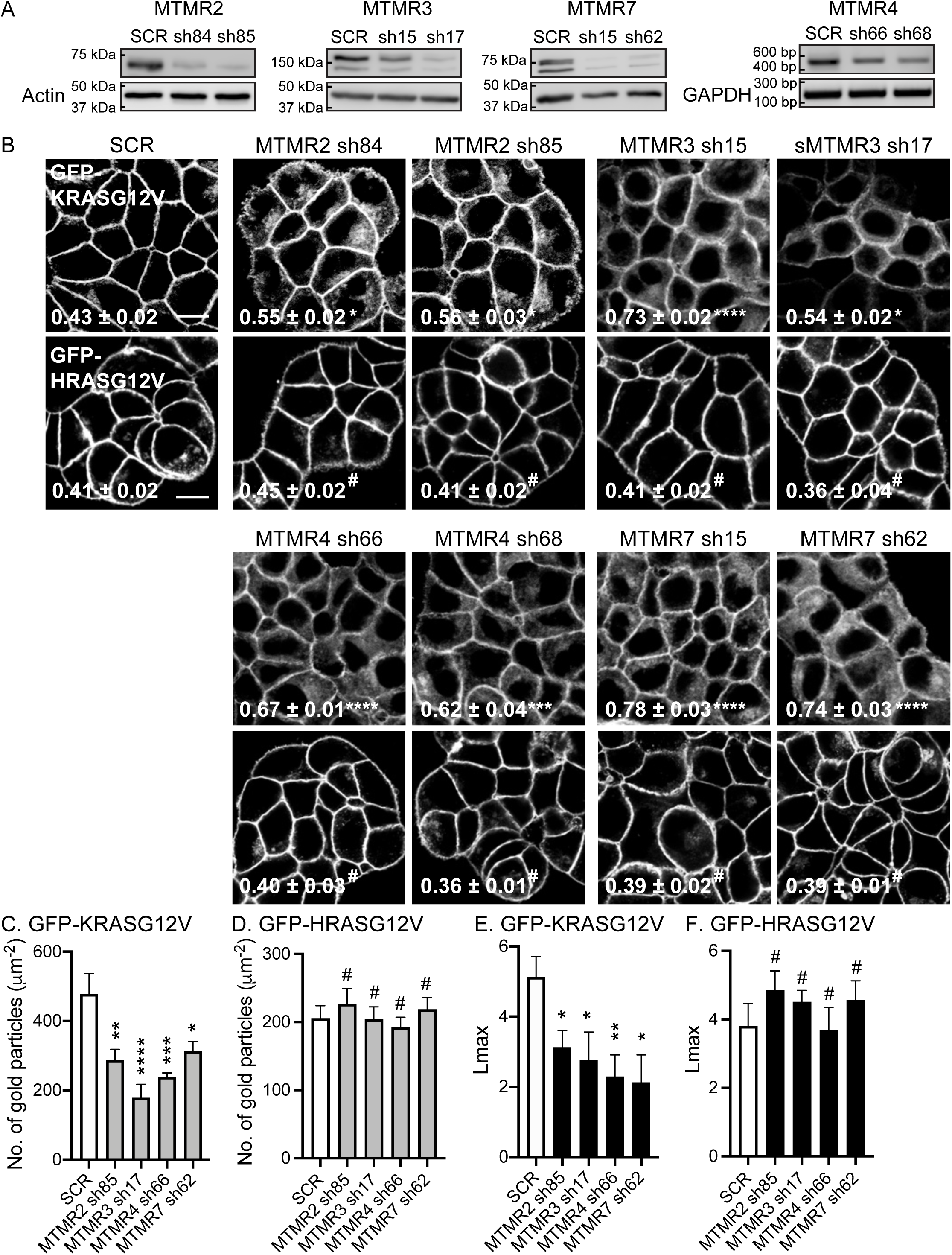
MTMR 2/3/4/7 regulate the PM localization of KRASG12V. (**A**) T47D cells stably expressing GFP-KRASG12V were infected with lentivirus expressing scrambled shRNA (SCR) or shRNA targeting *MTMR2, 3, 4* or *7*, followed by 1 ug/mL puromycin selection. Cell lysates were prepared and immunoblotted with anti-MTMR2, 3 and 7 antibodies. Actin blots are shown as loading controls. For MTMR4, cDNA was amplified with primers specific for *MTMR4* exons 1 and 5, or *GAPDH* exons 2 and 3 as a loading control. (**B**) T47D cells co-expressing shRNA targeting *MTMR 2/3/4/7* with GFP-KRASG12V or -HRASG12V were incubated with CellMask for 1 h at 37 °C incubator, fixed with 4% PFA and imaged by confocal microscopy. Representative GFP-KRASG12V and -HRASG12V images are shown. The corresponding CellMask and merged images for GFP-KRASG12V are shown in Fig. S1A. The inserted values represent a mean fraction ± S.E.M. of CellMask colocalizing with GFP-KRASG12V calculated by Manders’ coefficient from three independent experiments. Scale bar – 10 μm. (**C** and **D**) Intact basal PM sheets prepared from T47D cells co-expressing GFP-KRASG12V or - HRASG12V and shRNA targeting *MTMR2, 3, 4* or *7* were labeled with anti-GFP-conjugated gold particles and visualized by EM. Representative EM images are shown in Fig. S1B. The graphs show a mean number of gold particles ± SEM (n ≥ 15). (**D** and **E**) Spatial mapping of the same gold-labeled PM sheets was performed. The peak values, *L_max_*, representing the extent of KRAS and HRAS spatial organization, are shown as bar graphs (n ≥ 15). Significant differences between control (SCR-expressing) and *MTMR*-silenced cells were assessed by one-way ANOVA tests for (B – D) and bootstrap tests for (E and F) (* p < 0.05, ** p < 0.01, *** p < 0.001, **** p < 0.0001).

### MTMR 2/3/4/7 regulate the cellular level and distribution of PtdSer

The PM interaction of KRAS, but not HRAS, is dependent on PM PtdSer content (Cho et al., 2012b; Cho et al., 2016b; Garrido et al., 2020; Miller et al., 2019; Tan et al., 2018; van der Hoeven et al., 2018; Zhou et al., 2017). To quantify PM PtdSer content, T47D cells stably co-expressing GFP-LactC2, a well-characterized PtdSer probe (Yeung et al., 2008), and shRNA targeting *MTMR 2/3/4/7* were incubated with CellMask and imaged by confocal microscopy. GFP-LactC2 was primarily localized to the PM in control cells but was redistributed in *MTMR 2/3/4/7* KD cells as reflected by higher Manders’ coefficient values (Fig. 2A). EM analysis of isolated intact PM sheets prepared from T47D cells expressing GFP-LactC2 and incubated with anti-GFP antibody-conjugated gold shows that gold particle labeling density of the inner basal PM was significantly reduced when cells co-express shRNAs for *MTMR 2/3/4/7,* indicating depletion of inner PM PtdSer (Fig. 2B). The loss of inner leaflet PtdSer was not due to flipping to the outer leaflet since Annexin V binding to the outer PM leaflet, as measured by flow cytometry, did not increase in *MTMR 2/3/4/7* KD cells (Fig. 2C). Finally, whole cell lipidomics revealed that *MTMR 2/3/4/7* KD resulted in a significant reduction in total cellular PtdSer content (Figs. 2D and E). Together, these results show that MTMR 2/3/4/7 potently regulates the PM and total levels of PtdSer; consistent with a mechanistic role for MTMR 2/3/4/7 in controlling PtdSer cellular distribution.

**Figure 2.**
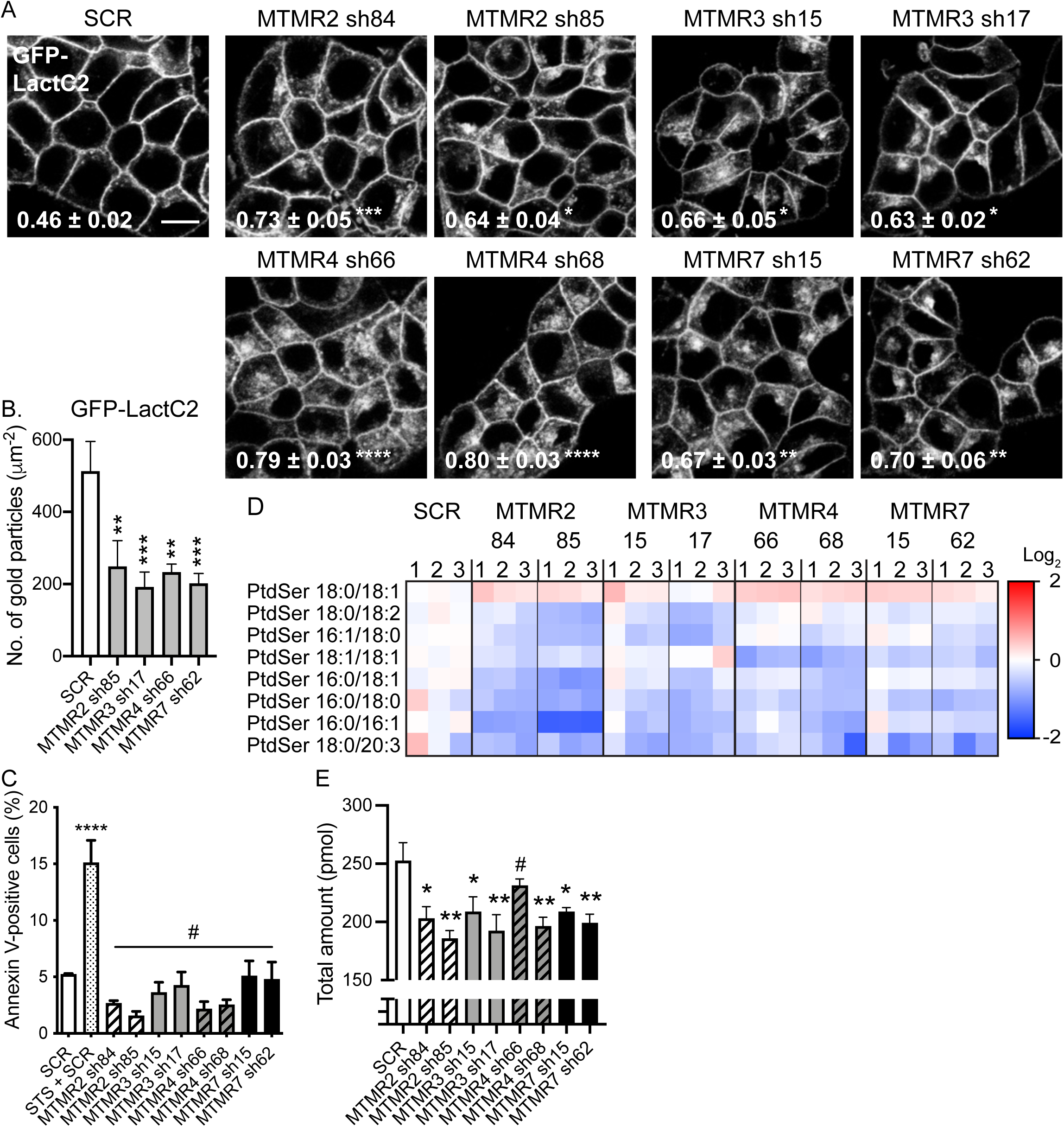
MTMR 2/3/4/7 regulate the cellular level and distribution of PtdSer. (**A**) T47D cells stably expressing GFP-LactC2 were infected with lentivirus expressing scrambled shRNA (SCR) or shRNA targeting *MTMR2, 3, 4* or *7*, followed by 1 ug/mL puromycin selection. Cells were incubated with CellMask for 1 h at 37 °C incubator, fixed with 4% PFA and imaged by confocal microscopy. Representative GFP-LactC2 images are shown. The inserted values represent a mean fraction ± S.E.M. of CellMask colocalizing with GFP-LactC2 calculated by Manders’ coefficient from three independent experiments. Scale bar – 10 μm. (**B**) Intact basal PM sheets prepared from T47D cells co-expressing GFP-LactC2 and shRNA targeting *MTMR2, 3, 4* or *7* were labeled with anti-GFP-conjugated gold particles and visualized by EM. The graphs show a mean number of gold particles ± SEM (n ≥ 15). (**C**) *MTMR 2/3/4/7* KD and stably expressing GFP-KRASG12V T47D cells were incubated with Cy5-conjugated annexin V and annexin V-positive cells were counted by flow cytometry. Scrambled (SCR) shRNA-expressing cells were treated with 1 μM staurosporine (STS) for 6 h to induce apoptosis. The graph represents a mean (%) ± S.E.M. of annexin V-positive cells from three independent experiments. (**D**) Whole cell PtdSer levels were measured in these cells via electron spray ionization and MS/MS from three independent experiments. A heat map was constructed to quantify the changes of different lipid species after *MTMR* knockdown. Each row represents a different species of PtdSer, while each column represents a single sample. The scaled expression values of each lipid measured is plotted in red-blue color log_2_ scale. In relation to control (SCR) cells, red and blue colored tiles indicate higher and lower lipid contents, respectively. (**E**) The graph shows the mean of total moles ± S.E.M. of PtdSer. Significant differences between control (SCR-expressing) and *MTMR*-silenced cells were assessed using one-way ANOVA tests (* p < 0.05, ** p < 0.01, *** p < 0.001, **** p < 0.0001, # – not significant).

### MTMR 2/3/4/7 regulate PI4P content at the PM

In mammalian cells, inner PM leaflet PI4P and PI(4,5)P_2_ recruit the LTPs, ORP5 and ORP8, to ER-PM membrane contact sites (MCSs) to exchange newly synthesized PtdSer in the ER for PI4P from the PM (Chung et al., 2015; Ghai et al., 2017; Moser von Filseck et al., 2015; Sohn et al., 2018). Depleting PM PI4P or PI(4,5)P_2_, concordantly depletes PtdSer and KRAS from the PM (Gulyas et al., 2017; Kattan et al., 2019). Depletion of PI4P at the Golgi complex also mislocalizes PtdSer and KRAS from the PM via an as yet unknown mechanism (Miller et al., 2019). To examine whether silencing *MTMR 2/3/4/7* perturbs cellular localization of PI4P and PI(4,5)P_2_, *MTMR 2/3/4/7* KD T47D cells expressing GFP-P4M-SidM or -PH-PLCδ1, probes for PI4P (Hammond et al., 2014) and PI(4,5)P_2_ (Varnai and Balla, 1998), respectively, were imaged by confocal microscopy. GFP-P4M-SidM decorated the PM and Golgi complex in control cells (closed and open arrowheads, respectively in Fig. 3A), consistent with previous studies (Hammond et al., 2014; Miller et al., 2019), whereas *MTMR 2/3/4/7* KD perturbed GFP-P4M-SidM staining of the PM, but not the Golgi complex (Fig. 3A). By contrast, the PM localization of GFP-PH-PLCδ1 was unchanged in *MTMR 2/3/4/7* KD cells (Fig. 3A). EM analysis of immunogold labelled PM sheets also shows significant reduction in PI4P PM content, but no change in PI(4,5)P_2_ PM content after *MTMR 2/3/4/7* KD (Fig. 3B and C). Given the reduction in PM PI4P content, we next examined ORP5 cellular localization. Confocal imaging showed that GFP-ORP5 predominantly localized to the ER-PM MCS in control cells, but substantially redistributed, presumably, to the ER, in *MTMR 2/3/4/7* KD cells (Fig. 3A) (Du et al., 2020; Monteiro-Cardoso et al., 2022). We conclude that MTMR 2/3/4/7 regulate ORP5 localization to the ER-PM MCS by maintaining the PM level of PI4P.

**Figure 3.**
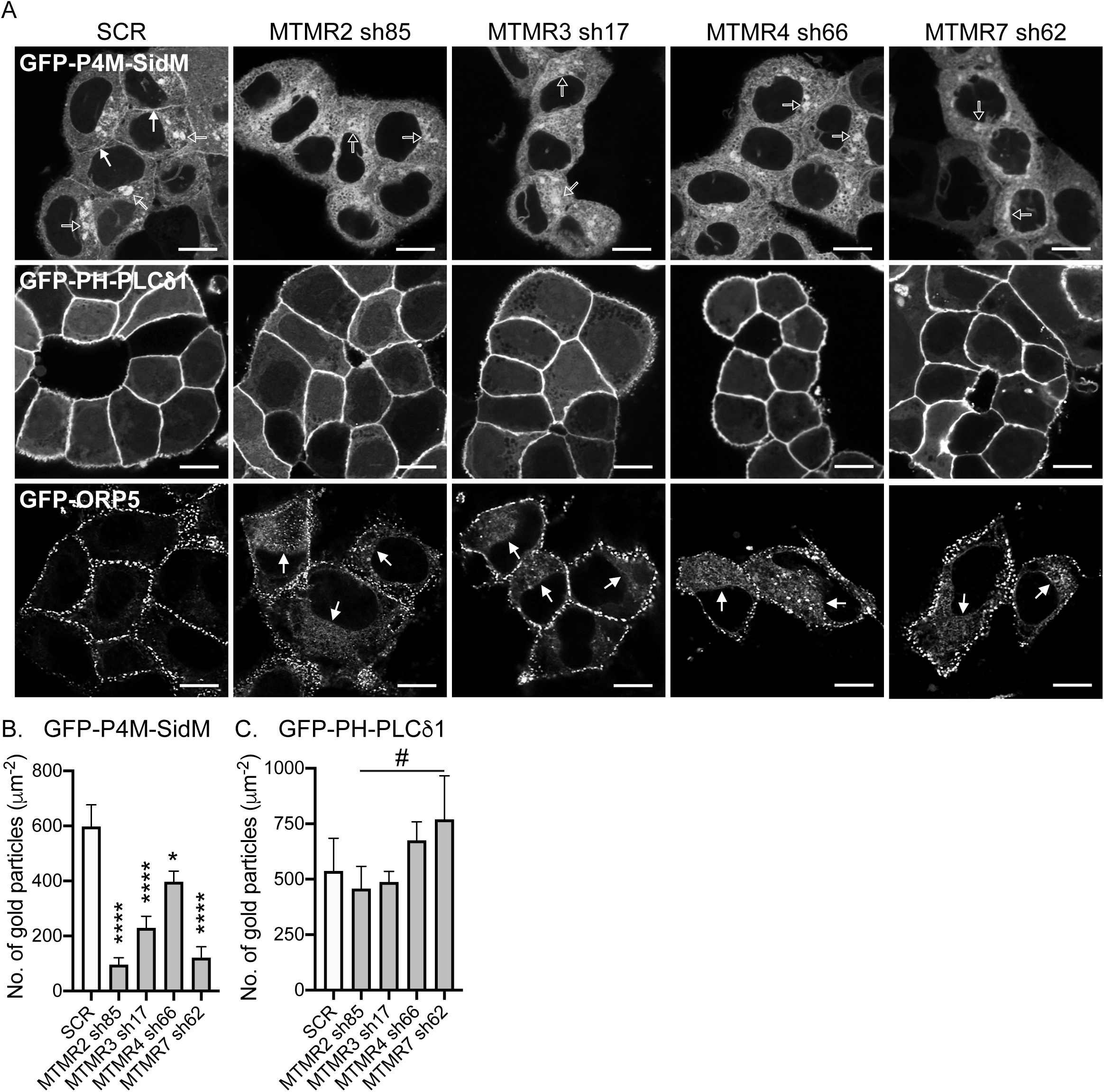
MTMR 2/3/4/7 regulate the localization of PM PI4P and ORP5. (**A**) WT T47D cells were infected with lentivirus expressing scrambled shRNA (SCR) or shRNA targeting *MTMR2, 3, 4* or *7*, followed by 1 ug/mL puromycin selection. These cells were transfected with GFP-P4M- SidM, -PH-PLCδ1, or -ORP5, and fixed with 4% PFA and imaged by confocal microscopy. Closed and open arrowheads for GFP-P4M-SidM indicate the staining of PM and Golgi complex, respectively. Closed arrowheads for GFP-ORP5 indicate the ER localization of ORP5. Scale bar – 10 μm. Intact basal PM sheet of Caco-2 cells co-expressing GFP-P4M-SidM (**B**) or -PH- PLCδ1 (**C**) with shRNA targeting *MTMR2, 3, 4,* or *7* were labeled with anti-GFP-conjugated gold particles and visualized by EM. The graphs show a mean number of gold particles ± SEM (n ≥ 15). Significant differences between control (SCR-expressing) and *MTMR*-silenced cells were assessed by one-way ANOVA tests (* p < 0.05, **** p < 0.0001, # – not significant).

### *MTMR 2/3/4/7* KD elevates PM PI3P content

PI3P is synthesized from PI by class II and III PI3Ks, whence it can be converted to PI(3,5)P_2_ or PI by PI 5-kinase, or myotubularin family phosphatases, respectively (Jean and Kiger, 2014; Pulido et al., 2013). To examine if depleting *MTMR 2/3/4/7* alters cellular PI3P levels or distribution, *MTMR 2/3/4/7* KD T47D cells expressing the well-characterized PI3P probe, GFP-2xFYVE, the tandem FYVE domain of Hrs (hepatocyte growth factor-regulated tyrosine kinase substrate) (Gillooly et al., 2000) were stained with CellMask and imaged by confocal microscopy. Manders’ coefficient values estimating fractions of endomembranes co-localized with GFP-2xFYVE-decorated endosomes were elevated in *MTMR 3/4/7*, but not *MTMR2*, KD cells, suggesting increased PI3P content in 2xFYVE-decorated endosomes (Figs. 4A and B). Also, individual KD of *MTMR 2/3/4/7* gene elevated total cellular PI3P level (Fig. 4C), consistent with previous studies (Cao et al., 2008; Lahiri et al., 2015; Ma et al., 2008; Zhao et al., 2019). Our data further show some PM decoration with GFP-2xFYVE in control and *MTMR* KD cells with no noticeable difference between cell lines, indicative of PI3P at the PM in these cell lines (arrowheads in Fig. 4A). To formally quantify PM PI3P content, we performed EM analysis on intact basal PM sheets prepared from *MTMR 2/3/4/7* KD T47D cells expressing GFP-2xFYVE. The data reveal that silencing *MTMR 2/3/4/7* significantly elevated PI3P levels in the PM compared to the control (Fig. 4D). Intriguingly, confocal and EM imaging further show GFP-MTMR 2/3/4/7 localization to the PM (Fig. 4E and F), supported by previous studies reporting the PM localization of other MTMR members, MTMR5 and 9 (Kim et al., 2003; Zou et al., 2009). We showed that while *MTMR 2/3/4/7* KD modestly increases PM PI3P level by less than 2-folds, the KD reduces PM PI4P, PtdSer and KRAS levels to greater extents by 2 to 3-folds (Figs. 1 – 3). Together, these data suggest that MTMR 2/3/4/7 localize to the PM, such that genetic depletion disrupts PI3P conversion to PI at the PM. This in turn, modestly increases total PI3P content including at the PM and 2xFYVE-decorated endosomes, which leads to substantial inhibition of the PM localization of KRAS, PtdSer and PI4P.

**Figure 4.**
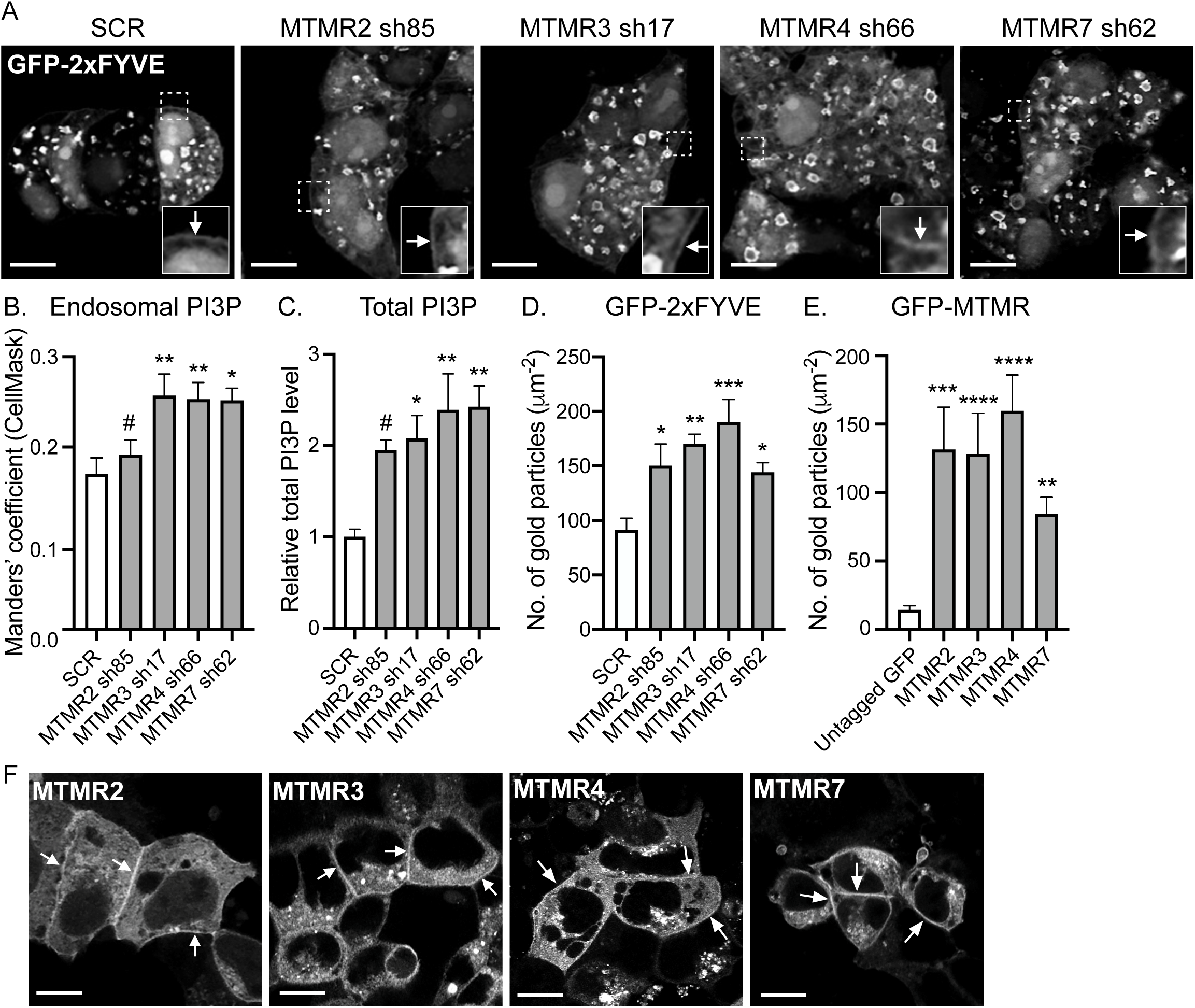
MTMR 2/3/4/7 regulate PM PI3P contents. (**A**) T47D cells expressing scrambled shRNA (SCR) or shRNA targeting *MTMR2, 3, 4* or *7*, followed by 1 ug/mL puromycin selection, were overexpressed with GFP-2xFYVE. Cells were incubated with CellMask for 1 h at 37 °C incubator, fixed with 4% PFA and imaged by confocal microscopy. Representative GFP- 2xFYVE images are shown. Their corresponding CellMask and merged images are shown in Fig. S3. A selected region (the white square) is shown at a higher magnification. Arrowheads indicate the PM-staining of GFP-2xFYVE. (**B**) The graph represents a mean fraction ± S.E.M (n = 13) of CellMask colocalizing with GFP-2xFYVE calculated by Manders’ coefficient. (**C**) Acidic lipids were extracted from *MTMR 2/3/4/7* KD T47D cells, and total PI3P was measured by a quantitative ELISA. The graph shows a relative mean total PI3P level ± S.E.M from three independent experiments. (**D**) Intact basal PM sheets prepared from these T47D cells from (A), or WT T47D cells expressing GFP-MTMR members (**E**) were labeled with anti-GFP-conjugated gold particles and visualized by EM. The graphs show a mean number of gold particles ± SEM (n ≥ 15). (**F**) Cells from (E) were fixed with 4% PFA and imaged by confocal microscopy. Arrowheads indicate the PM localization of GFP-MTMR members. Scale bar – 10 μm. Significant differences between control (SCR-expressing) and *MTMR*-silenced cells were assessed by one-way ANOVA tests for (B – D) (* p < 0.05, ** p < 0.01, *** p < 0.001, **** p < 0.0001, # - not significant). Significant differences between untagged GFP (control) and GFP- MTMR-expressing cells were assessed by one-way ANOVA test for (E) (** p < 0.01, *** p < 0.001, **** p < 0.0001).

### The enzymatic activity of MTMR regulates the PM localization of KRAS, PtdSer and PI4P

To examine if MTMR requires its enzymatic activity for maintaining the PM localization of KRAS, PtdSer and PI4P, we overexpressed mCherry-tagged MTMR3 WT or C413S, the phosphatase-inactive mutant (Walker et al., 2001), in *MTMR3* KD T47D cells stably expressing GFP- KRASG12V or -LactC2, and analyzed their cellular localization by confocal and EM. Our data show that while overexpression of MTMR3 restored the PM binding of GFP-KRASG12V and - LactC2, the C413S mutant did not show any changes (Figs. 5A – D). We further assessed the PM PI4P and PI3P contents by measuring the PM binding of GFP-P4M-SidM and -2xFYVE, respectively, by EM. Like KRASG12V and PtdSer, MTMR3 WT overexpression promoted the PM binding of GFP-P4M-SidM and -2xFYVE, but not the C413S mutant (Figs. 5E – F). We repeated the experiments with MTMR4 WT and the C407S, the phosphatase-inactive mutant (Lorenzo et al., 2006), and observed a similar trend. Overexpression of WT MTMR4, but not the inactive mutant, reduced PM PI3P levels, and promoted the PM binding of GFP-KRASG12V, - LactC2 and -P4M-SidM (Fig. S4). Moreover, despite the similar expression level, ectopic expression of WT MTMR 2/4/7 did not restore the effects of *MTMR3* KD (Figs. 5A – G). We made a similar observation with rescuing *MTMR4* KD, in which overexpression of WT MTMR 2/3/7 did not revert the effects of *MTMR4* KD (Fig. S4). Together, these data suggest that MTMR members require their enzymatic activity for maintaining the PM localization of KRAS, PtdSer and PI4P, and that the enzymatic activity of each MTMR is not functionally redundant.

**Figure 5.**
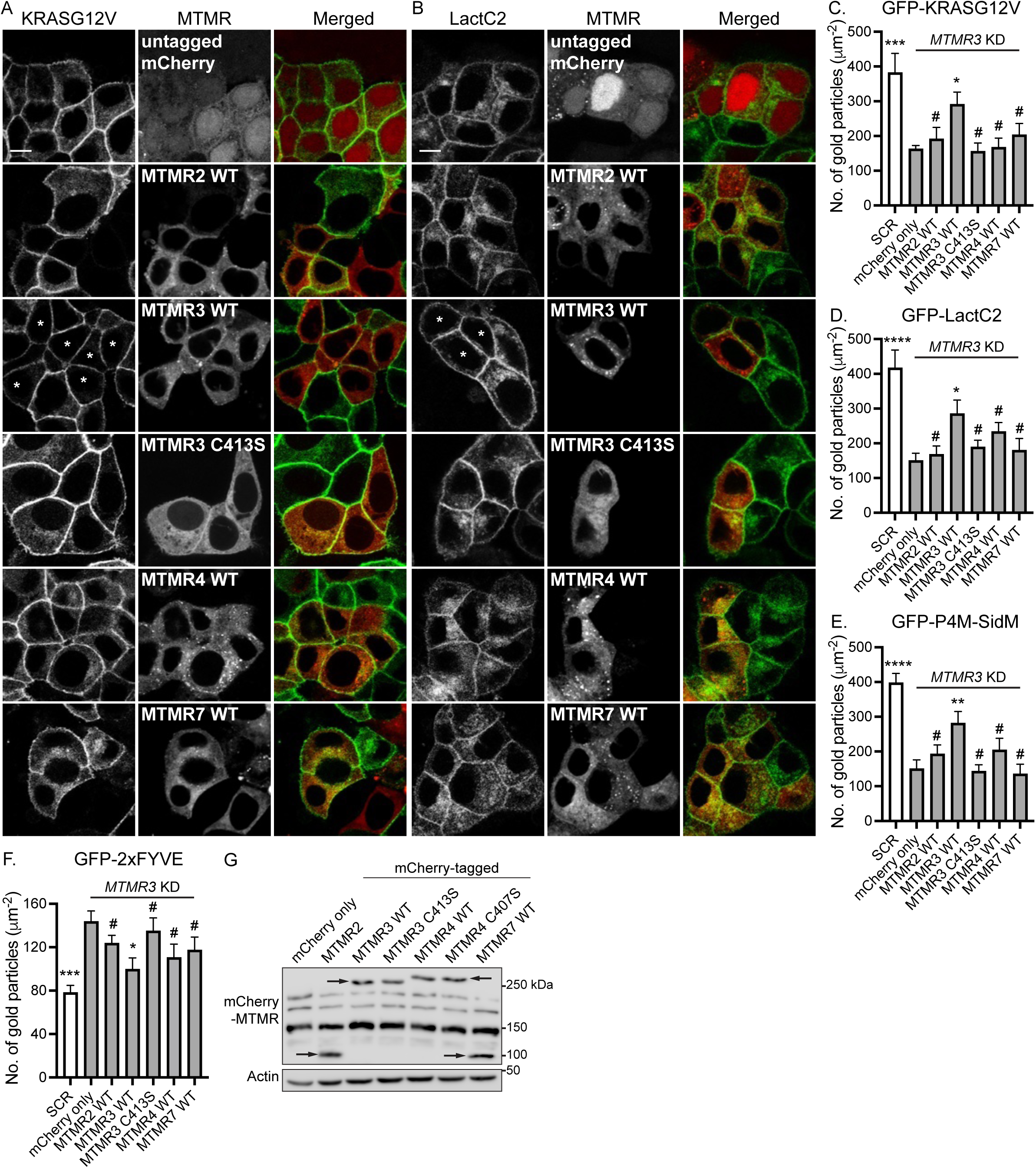
Ectopic expression of WT MTMR3 restores the effects of *MTMR3* loss. T47D cells co-expressing *MTMR3* shRNA17 and GFP-KRASG12V (**A**) or -LactC2 (**B**) were infected with lentivirus expressing indicated mCherry-MTMR proteins, fixed with 4% PFA and imaged by confocal microscopy. Scale bar – 10 μm. Asterisks indicate the restored PM localization of GFP- KRASG12V or -LactC2. Intact basal PM sheets were prepared from T47D cells co-expressing *MTMR3* shRNA17 and GFP-KRASG12V (**C**) or -LactC2 (**D**), -P4M-SidM (**E**), or -2xFYVE (**F**) were infected with lentivirus expressing indicated mCherry-MTMR members, and labeled with anti-GFP-conjugated gold particles and visualized by EM. The graphs show a mean number of gold particles ± S.E.M. (n = 10). (**G**) Cell lysates from T47D cells co-expressing GFP- KRASG12V, *MTMR3* shRNA17 and mCherry-MTMR members were immunoblotted with anti- RFP antibody. Arrowheads represents the expression of indicated mCherry-MTMR members. An actin blot is shown as a loading control. Representative blots are shown from three independent experiments. Significant differences between control (*MTMR3* KD and untagged mCherry-expressing) and other cells were assessed by one-way ANOVA tests for (C – F) (* p < 0.05, ** p < 0.01, *** p < 0.001, **** p < 0.0001, # - not significant).

### Protein expression of MTMR 2/3/4/7 is interdependent

While there are 9 catalytically active MTMR members (Wang et al., 2024), single KD of *MTMR 2/3/4/7* is sufficient to elevate PI3P contents and dissociates KRAS, PtdSer and PI4P from the PM (Figs. 1 – 4). One plausible explanation is that depleting single MTMR member may perturb the enzymatic activities of other MTMR members, resulting in significant disruption of PI3P metabolism. To test this notion, we first examined if the loss of multiple MTMR members can induce additive effects in these phenotypes. *MTMR3* KD T47D cells were infected with lentivirus expressing shRNA targeting *MTMR2, 4,* or *7*, but the double KD resulted in cytotoxicity (not shown). We then measured endogenous protein levels of MTMR 2/3/7 after KD of single MTMR member. Our immunoblot data show that the protein expression of MTMR 2/3/7 were concomitantly reduced upon *MTMR2*, *3*, *4* or *7* KD (Fig. 6A), suggesting the protein expression of MTMR 2/3/7 are interdependent. Moreover, while ectopic expression of WT MTMR3 in *MTMR3* KD cells restored the protein levels of MTMR2 and 7, overexpression of WT MTMR2, 4 or 7 in *MTMR3* KD cells did not restore the endogenous MTMR3 protein level (Fig. 6B and C). These data suggest that the protein expression, and thereby the enzymatic activity, of MTMR 2/3/4/7 are interdependent.

**Figure 6.**
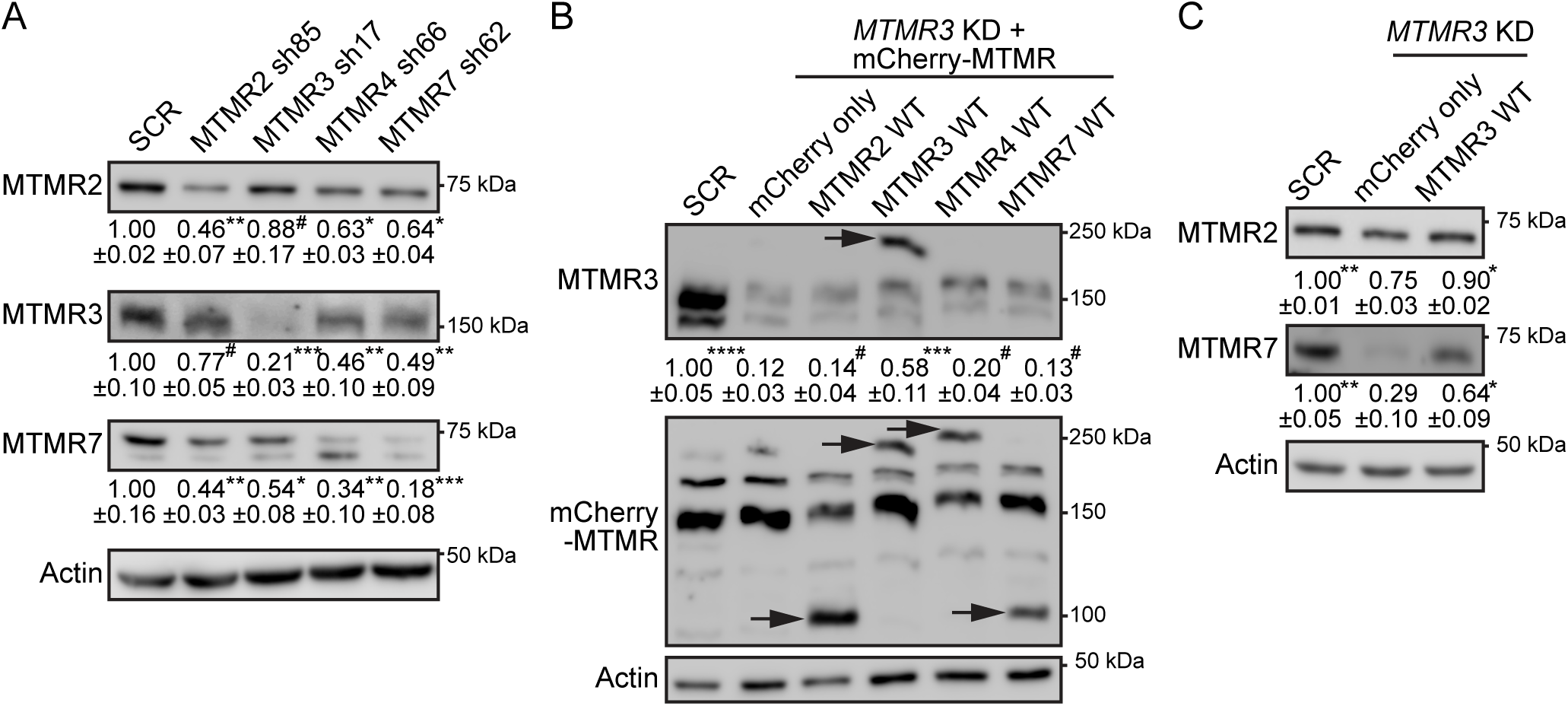
Protein expression of MTMR 2/3/4/7 is interdependent. (**A**) Cell lysates from T47D cells co-expressing GFP-KRASG12V and indicated *MTMR*-targeting shRNA were immunoblotted for endogenous MTMR2, 3 and 7. (**B**) Cell lysates from T47D cells co-expressing GFP-KRASG12V and *MTMR3* shRNA17, and infected with lentivirus expressing indicated mCherry-MTMR members were prepared and immunoblotted for endogenous MTMR3. The membrane was then stripped and re-blotted with anti-RFP antibody. Arrowheads represent the expression of indicated mCherry-tagged MTMR members. (**C**) Cell lysates from (B) were blotted for endogenous MTMR2 and 7. Actin blots are shown as loading controls. Representative blots are shown. Values indicate mean MTMR member ± S.E.M relative to SCR-expressing cells from three independent experiments. Significant differences between control (*MTMR3* KD and untagged mCherry-expressing) and other cells were assessed by one-way ANOVA tests (* p < 0.05, ** p < 0.01, *** p < 0.001, **** p < 0.0001, # - not significant).

### MTMR 2/3/4/7 regulate KRAS signaling

Since *MTMR 2/3/4/7* KD dissociates KRASG12V from the PM, we used two biological systems to evaluate effects on KRAS signal transmission. First, we analyzed signaling by the KRAS ortholog, let-60 in the invertebrate model system *Caenorhabditis elegans* (*C. elegans*), where constitutively active mutation of *let-60* induces a quantifiable multivulva phenotype (Reiner et al., 2008). We used RNAi designed to target *C. elegans* MTMR orthologs and examined whether *MTMR* KD could suppress the multivulva phenotype associated with expression of LET-60 G13D (n1046). Our data show that silencing *mtm-3* (ortholog of *MTMR3*), or *mtm-6* (ortholog of *MTMR7*) abrogated multivulva formation, and indeed were equipotent with two previously reported potent suppressors of LET-60 G13D signaling (*heo-1* and *riok-1*) (Smith and Levitan, 2004; Weinberg et al., 2014) (Fig. 7A). Secondly, we examined it in human cancer model cell lines. The proliferation of *MTMR 2/3/4/7* KD T47D cells stably expressing GFP-KRASG12V, but not -HRASG12V, was significantly inhibited compared to parental cells replete for each *MTMR* gene (Fig. 7B and C), suggesting the growth inhibitory effects of *MTMR 2/3/4/7* KD is KRAS-specific. We also examined the proliferation and RAS signaling in human pancreatic ductal adenocarcinoma (PDAC) cell lines, MIA PaCa-2 and PANC-1, which harbor KRAS with activating point mutation, G12C and G12D, respectively, and their growth depends on the oncogenic KRAS signaling (Hayes et al., 2016; Kovar et al., 2020; Rehl et al., 2023). For a control PDAC cell line, we included BxPC-3, which harbors WT KRAS, and its growth is independent of the KRAS signaling. We showed that *MTMR 2/3/4/7* KD inhibited the proliferation of MIA PaCa-2 and PANC-1, but not BxPC-3 cells (Fig. 7D). For assessing RAS signal output, we measured active phosphorylation sites of ERK1/2 and Akt for the RAS/MAPK and RAS/PI3P/Akt pathways, respectively. Our immunoblotting data show a tendency of decreased phospho-ERK after *MTMR 2/3/4/7* KD in KRAS-dependent PDAC (Fig. 7E). To further examine the ERK activity, we measured the expression level of the transcriptional factor, c-MYC, since active ERK enhances its stability (Sears et al., 1999). Our data show that c-MYC protein level was significantly decreased upon *MTMR 2/3/4/7* KD in KRAS-dependent, but not -independent PDAC cell lines, indicative of blocked MAPK signaling in KRAS-dependent PDAC (Fig. 7E). Akt phosphorylation was elevated in *MTMR2* and *7* KD cells, while no significant changes were observed upon *MTMR3* and *4* KD in all three cell lines, suggesting that MTMR 2/3/4/7 differentially regulate the PI3K/Akt signaling in an oncogenic KRAS-independent manner. We further measured endogenous KRAS expression level since KRAS protein stability is dysregulated after the PM dissociation (Cho et al., 2012b; Garrido et al., 2020; Kovar et al., 2020). Our immunoblot data show that endogenous KRAS expression was significantly reduced upon *MTMR 2/3/4/7* KD in KRAS-dependent, but not -independent PDAC cell lines (Fig. 7E). Together, our study provides strong evidence that the MTMR phosphatases regulate KRAS signal transmission by maintaining PtdSer and KRAS PM localization and their genetical loss dissociates the PM localization of PtdSer and KRAS, but not HRAS, resulting in the inhibition of KRAS/MAPK/c-MYC signal output and instability of KRAS protein expression in KRAS-dependent PDAC.

**Figure 7.**
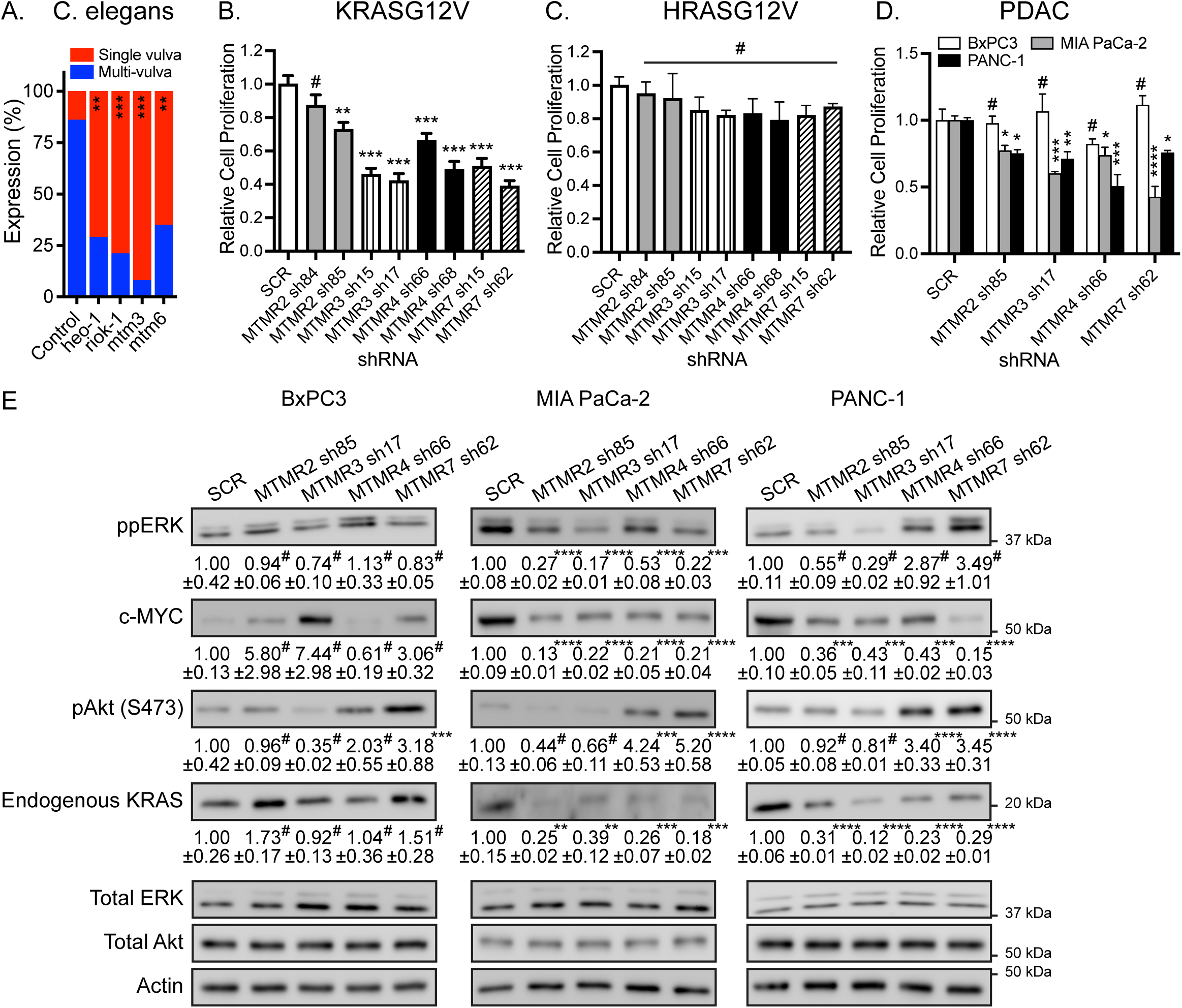
MTMR 2/3/4/7 regulate the oncogenic KRAS signal output. (**A**) RNAi was induced by feeding *let-60* (n1046) L1 worms through adult stage with *E. coli* strain HT115, producing dsRNA to target genes. The presence of the multi-vulva phenotype was scored using DIC/Nomarski microscopy. 100 – 200 worms were assayed per RNAi knockdown. T47D cells stably expressing (**B**) GFP-KRASG12V or (**C**) -HRASG12V, or (**D**) WT BxPC3, MIA PaCa-2 and PANC-1 cells were infected with lentivirus expressing scrambled shRNA (SCR) or shRNA targeting *MTMR 2/3/4/7*, followed by 1 μg/mL puromycin selection. Cells were seeded on a 96- well plate and cultured for 4 days. Complete growth medium was replaced every 24 h, and cell numbers were counted to measure cell proliferation. Graphs show the mean cell proliferation ± S.E.M. relative to that for the control cells (SCR-expressing) from three independent experiments. (**E**) Cell lysates prepared from (D) were immunoblotted for ppERK, pAkt (S473), c- MYC and endogenous KRAS. Values indicate the mean ppERK, pAkt, c-MYC or endogenous KRAS ± S.E.M relative to the control cells (SCR-expressing) from three independent experiments. Total ERK, Akt and actin blots were used as loading controls. Representative blots are shown. Significant differences between control (SCR-expressing) and *MTMR* KD cells were assessed using one-way ANOVA tests (* p<0.05, ** p<0.01, *** p<0.001, # - not significant).

## Discussion

We have identified new roles for the phosphatases, MTMR 2/3/4/7, in maintaining the PM localization of PI4P, PtdSer and KRAS. We show that KD of *MTMR 2/3/4/7* dissociates PtdSer and KRAS from the PM, and also reduces total cellular PtdSer level. Concomitantly, *MTMR 2/3/4/7* KD elevates PM PI3P content, reduces PM PI4P content and dissociates ORP5 from the ER-PM MCS. Collecting these observations together, we propose that the PM-localized MTMR phosphatases are required to help maintain PM levels of PI for generating PM PI4P by PI4KA, which in turn maintains total PtdSer levels and drives ER to PM transport of PtdSer. PI is synthesized from phosphatidic acid at the cytosolic face of ER, where a fraction of PI is used for the synthesis of glycosylphosphatidylinositol-anchored proteins (Blunsom and Cockcroft, 2020), with the remaining PI being widely distributed across intracellular cytosolic membranes (Ashlin et al., 2021; Pemberton et al., 2020; Zewe et al., 2020). PI delivered from the ER directly to the PM by the LTPs, Nir2/3 and TMEM24 (Posor et al., 2022), is a crucial substrate for PM polyphosphoinositides (PPIn). Simple overexpression of Nir2/3 and TMEM24 elevates the synthesis and total amount of PM PPIn (Chang et al., 2013; Kim et al., 2015; Lees et al., 2017), suggesting that PM PI is constantly converted to PPIn and the PI supply to the PM is rate-limiting. In this context, the pathway to PI generation from PI3P by MTMR 2/3/4/7 at the PM, emerges as a critical alternative supplier of PI to allow sufficient PM PI4P generation hence PtdSer transport by the ORP5/8 machinery (Fig. 8). In addition to the PM, MTMR 2/3/4/7 localize to endosomes and promotes their trafficking by regulating endosomal PI3P and PI(3,5)P_2_ levels (Bonneick et al., 2005; Franklin et al., 2011; Mochizuki and Majerus, 2003; Naughtin et al., 2010; Previtali et al., 2007; Walker et al., 2001). Thus, perturbing endosomal trafficking by *MTMR 2/3/4/7* KD will block the delivery of endosomal PI to the PM, further contributing to the reduced PM PI content. Therefore, when *MTMR 2/3/4/7* is depleted, PM PtdSer level falls and KRAS dissociates from the PM (Fig. 8). We did not investigate the mechanism whereby *MTMR 2/3/4/7* KD reduced total cellular PtdSer content, but previous work has shown that PM PI4P depletion indirectly blocks PtdSer synthase 1 and 2 activities (Sohn et al., 2016). Since PM PI4P levels were reduced in *MTMR 2/3/4/7* KD cells, the same mechanism is likely responsible. Furthermore, inactivating mutations in *MTMR2* are found in Charcot-Marie-Tooth disease type 4B1, a severe autosomal recessive neuropathy with demyelination and myelin outfoldings of the nerve. Studies have proposed that disrupted endosomal trafficking by elevating endosomal PI3P and PI(3,5)P_2_ due to inactive MTMR2 likely contributes to the demyelination of nerve cells (Bonneick et al., 2005; Previtali et al., 2007). Intriguingly, PtdSer is the main component of myelin sheath (Ma et al., 2022), and inhibition of PI4KA in Schwann cells reduces PM and total levels of PtdSer and PI4P, and induces aberrant myelination (Alvarez-Prats et al., 2018). Thus, in addition to disrupted endosomal trafficking by *MTMR2* inactivation, it is plausible that the reduced PM levels of PtdSer and PI4P further contributes to the demyelination observed in Charcot-Marie-Tooth disease type 4B1.

**Figure 8.**
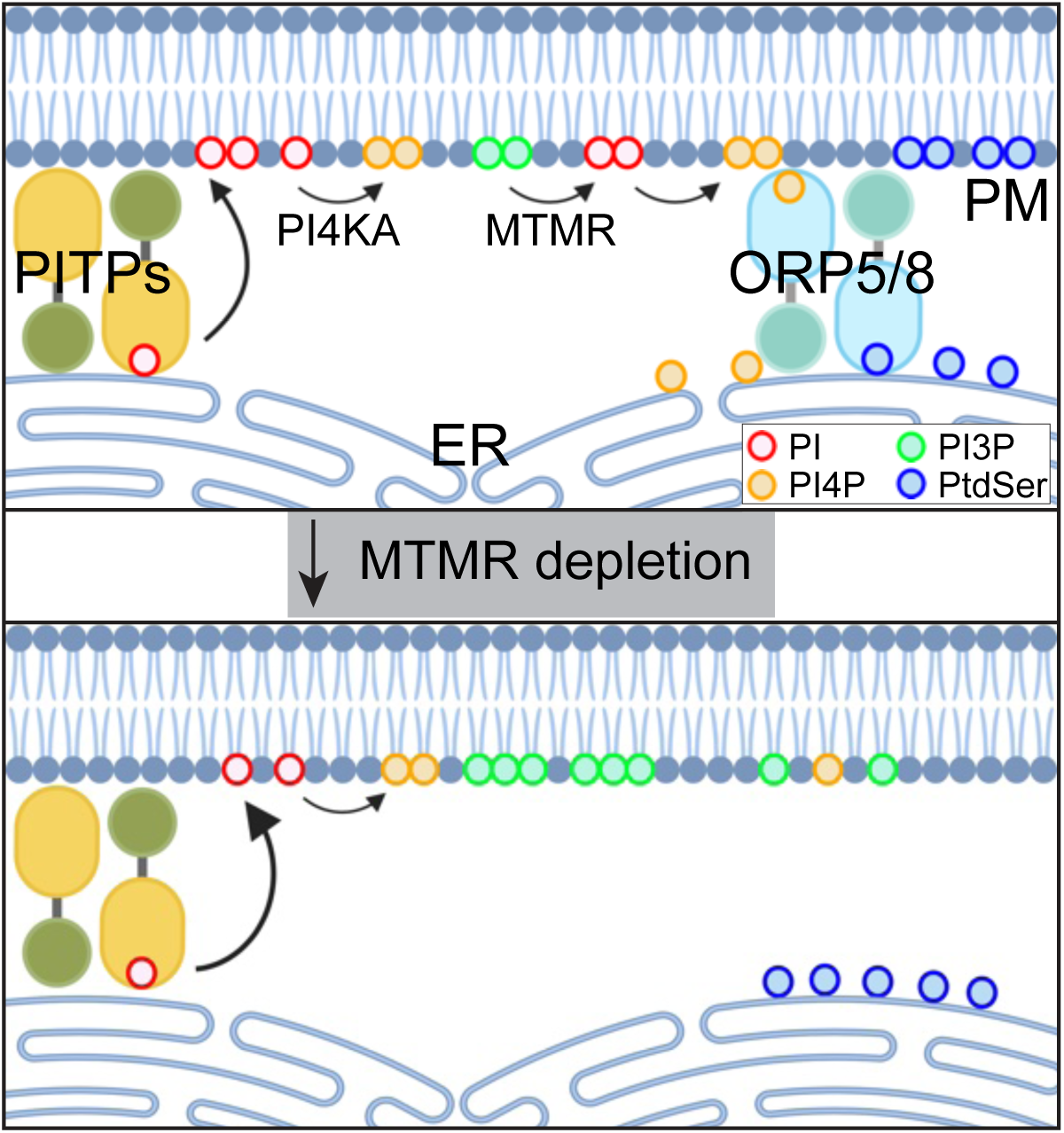
A working model of our study. PITPs deliver newly synthesized PI from the ER to PM, and MTMR at the PM concomitantly converts PI3P to PI, which are which are crucial substrate for PM PPIn like PI4P. ORP5 and 8, the lipid transfer proteins for PtdSer, exchanges ER PtdSer with PM PI4P at ER-PM membrane-contacting sites, which enriches PM PtdSer contents. This in turn, allows stable PM binding of KRAS, and thereby KRAS signal output. When MTMR is depleted, PM PI3P conversion to PI is blocked, providing insufficient PM PI contents for PI4P synthesis. This dissociates ORP5/8 from ER-PM membrane-contacting sites, resulting in reduced PM localization of PtdSer and thereby KRAS. PITPs – PI transfer proteins, PI4KA – phosphatidylinositol 4-kinase IIIα, ORP – oxysterol-binding protein related protein, PM – plasma membrane, ER – endoplasmic reticulum.

We have demonstrated that the protein expression of MTMR 2/3/4/7 are interdependent, whence single KD of *MTMR2, 3, 4* or *7* concomitantly reduces the protein levels of MTMR 2/3/4/7 (Fig. 6A), which likely accounts for the observation that a loss of *MTMR2, 3, 4 or 7* is sufficient to elevate cellular PI3P level despite the presence of 9 catalytically active MTMR members (Cao et al., 2008; Lahiri et al., 2015; Ma et al., 2008; Zhao et al., 2019). Also, while overexpression of WT MTMR3 in *MTMR3* KD cells restores endogenous protein levels of MTMR 2/4/7, overexpression of WT MTMR 2/4/7 does not restore endogenous MTMR3 protein level (Fig. 6B and C). These data are consistent with our rescue experiments, in which overexpression of WT MTMR3, but not the other MTMR members, in *MTMR3* KD cells restores the KD effects of *MTMR3*. For the interdependence of MTMR 2/3/4/7 protein expression, one plausible model is that the catalytically active and inactive MTMR members form one large, interconnected machinery, which may regulate the subcellular localization of MTMR members for maintaining cellular PI3P metabolism. The 9 enzymatically active MTMR members need to form homo- or heterodimers with the 7 inactive members via its coiled-coil domain for their enzymatic activity and subcellular localization (Hnia et al., 2012; Wang et al., 2024). MTMR3 and 4 form homodimers and heterodimers with each other (Lorenzo et al., 2006). MTMR2 form homodimers and heterodimers with inactive members, MTMR5, 12 and 13 (Kim et al., 2003; Nandurkar et al., 2003; Robinson and Dixon, 2005), while MTMR7 heterodimerizes with the inactive MTMR9 (Mochizuki and Majerus, 2003). Also, MTMR2 promotes MTMR13 membrane localization and they depend on each other for maintaining their protein expression (Ng et al., 2013; Robinson and Dixon, 2005). Thus, the loss of a single MTMR member could perturb the subcellular localization of its partner MTMR members, resulting in their protein degradation. This in turn, disrupts the interconnected network of MTMR members, resulting in the subcellular mislocalization and degradation of multiple MTMR members, and thereby perturbed PI3P metabolism. A systematic analysis of subcellular localization and protein expression of catalytically active and inactive MTMR members after genetical depletion of single MTMR member will provide deeper insights into the interdependence of MTMR members.

We have observed that *MTMR 2/3/4/7* KD reduces oncogenic KRAS protein expression (Fig. 7E). Previous studies have reported that *MTMR* KD stimulates autophagy via elevating PI3P content in the phagophore, a double-membrane structure that matures into autophagosome, which recruits autophagy-related proteins for autophagosome biogenesis (Nishimura and Tooze, 2020; Wang et al., 2024). Also, KRAS degradation occurs via the autophagy/lysosome-mediated mechanism, in which *ADAM9* loss or inhibition enhances the interaction between KRAS and plasminogen activator inhibitor 1 (PAI-1), which acts as a specific autophagy receptor binding to light chain 3 (LC3) for stimulating lysosomal degradation of KRAS (Huang et al., 2024). Together, our data suggest that the PM dissociated KRAS undergoes its degradation by enhanced autophagy induced by *MTMR* KD. However, this may only partially account for our observation. Since *MTMR 2/3/4/7* KD induces PM dissociation of KRAS without the G-domain (Fig. S2), it is likely that the KD induces the PM dissociation of WT KRAS expressed in BxPC-3, which was not degraded upon *MTMR 2/3/4/7* KD (Fig. 7E). Thus, the KRAS degradation upon *MTMR 2/3/4/7* KD is unlikely induced by KRAS PM dissociation. Also, while oncogenic KRAS/MAPK signaling promotes autophagy in PDAC (Bryant et al., 2019), we observed *MTMR 2/3/4/7* KD reduces the MAPK/c-MYC signaling in PDAC harboring oncogenic KRAS. Thus, the inhibition of MAPK/c-MYC signaling by *MTMR* KD would block autophagy, but promote it. Deeper insights for KRAS degradation in terms of oncogenic KRAS signaling and cellular PI3P content need to be elucidated.

In sum, our study proposes a new role for the MTMR phosphatases in maintaining PM PI4P and PtdSer contents by maintaining an adequate supply of PM PI from PI3P. Depleting MTMR 2/3/4/7 in turn, reduces PM and total cellular PtdSer levels, blocking KRAS PM localization and KRAS signaling. Thus, the mechanisms that maintain PM PI content may contain useful targets to abrogate oncogenic KRAS signaling. Our study further proposes an interdependent network of MTMR members for maintaining their protein expression and their phosphatase activity.

## Acknowledgements

This work was supported by Wright State University Seed Grant and the National Cancer Institute [R00-CA188593 to K.-J.C.], National Institute of General Medical Sciences [2R35GM119412 to G.R.V. Hammond] and Cancer Prevention and Research Institute of Texas [CPRIT RP200047 to JFH]. We would like to thank Karen Henkels for her contribution to this work.

## Authors contribution and ORCID IDs

- Taylor E. Lange performed investigation and validation.
- Ali Naji () performed investigation and validation.
- Ransome van der Hoeven () performed investigation data analysis.
- Hong liang performed investigation.
- Yong Zhou performed investigation, validation and data analysis.
- Gerald R.V. Hammond performed editing and review the manuscript.
- John F. Hancock performed funding acquisition and edit/review the manuscript.
- Kwang-jin Cho performed conceptualization, data curation, formal analysis, funding acquisition, investigation, methodology, project administration, resources, supervision, validation, visualization, and writing the draft and editing.

---

**Supplement Figure 1.**
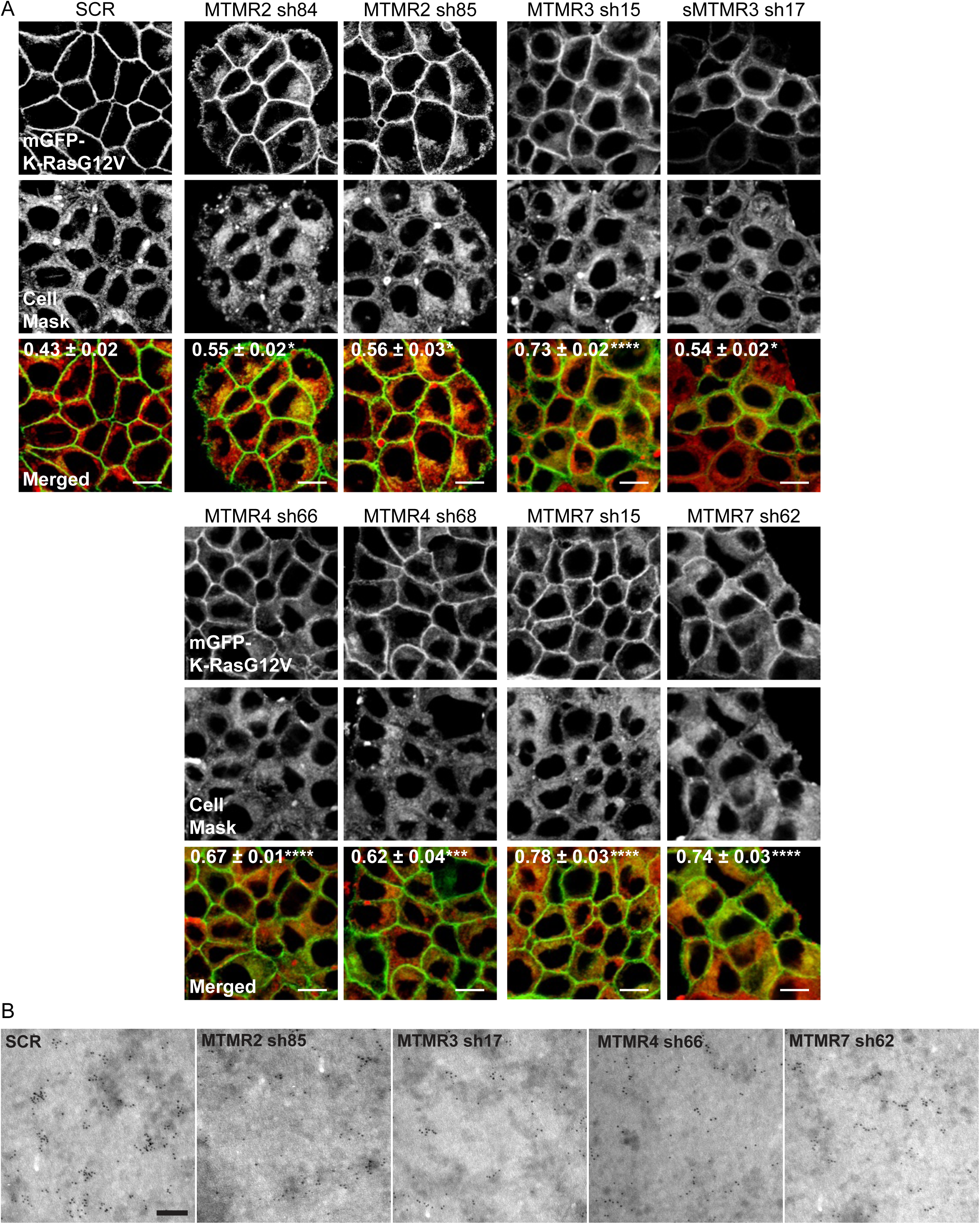
MTMR 2/3/4/7 regulate the PM localization of KRASG12V. (**A**) T47D cells stably expressing GFP-KRASG12V were infected with lentivirus expressing scrambled shRNA (SCR) or shRNA targeting *MTMR2, 3, 4* or *7*, followed by 1 ug/mL puromycin selection. Cells were incubated with CellMask for 1 h at 37 °C incubator, fixed with 4% PFA and imaged by confocal microscopy. The inserted values represent a mean fraction ± S.E.M. of CellMask colocalizing with GFP-KRASG12V calculated by Manders’ coefficient from three independent experiments. Scale bar – 10 μm. (**B**) Intact basal PM sheets prepared from T47D cells co-expressing GFP-KRASG12V and shRNA targeting *MTMR2, 3, 4* or *7* were labeled with anti-GFP-conjugated gold particles and visualized by EM. Representative EM images are shown. Scale bar – 0.1 μm. Significant differences between control (SCR-expressing) and *MTMR*- silenced cells were assessed by one-way ANOVA tests (* p < 0.05, *** p < 0.001, **** p < 0.0001).

**Supplement Figure 2.**
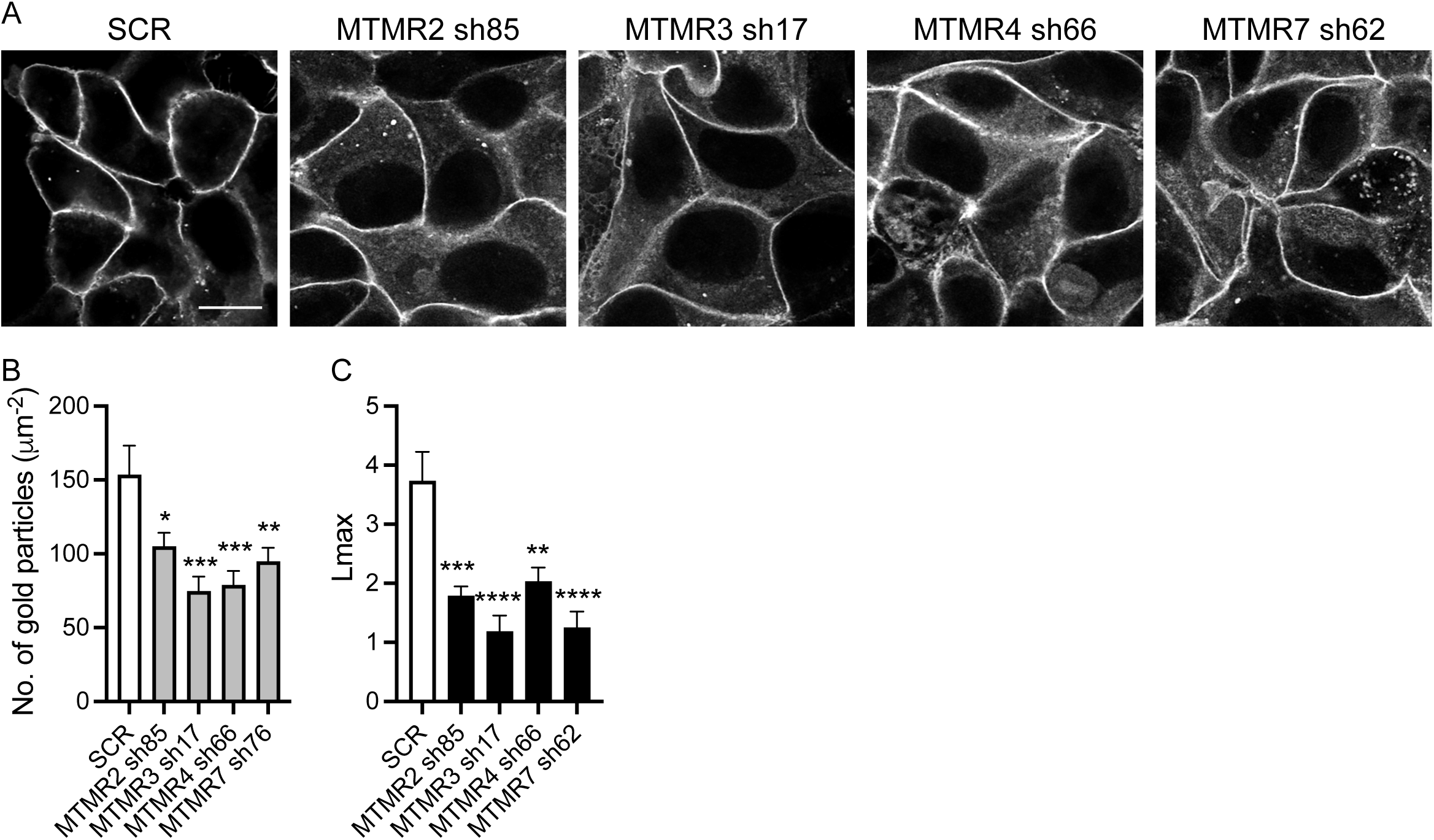
MT*MR 2/3/4/7* KD disrupts the PM localization and nanoclustering of CTK. (**A**) MCF-7 cells stably co-expressing GFP-CTK (the C-terminal membrane anchor of KRAS without the G-domain) and shRNA targeting MT*MR2, 3, 4 or 7* were fixed with 4% PFA and imaged by confocal microscopy. (**B**) Intact basal PM sheets of cells from (A) were prepared, labeled with anti-GFP-conjugated gold particles and visualized by EM. Scale bar – 10 μm. Significant differences between control (SCR-expressing) and *MTMR*-silenced cells were assessed by one-way ANOVA tests (* p < 0.05, ** p< 0.01, *** p < 0.001, **** p < 0.0001).

**Supplement Figure 3.**
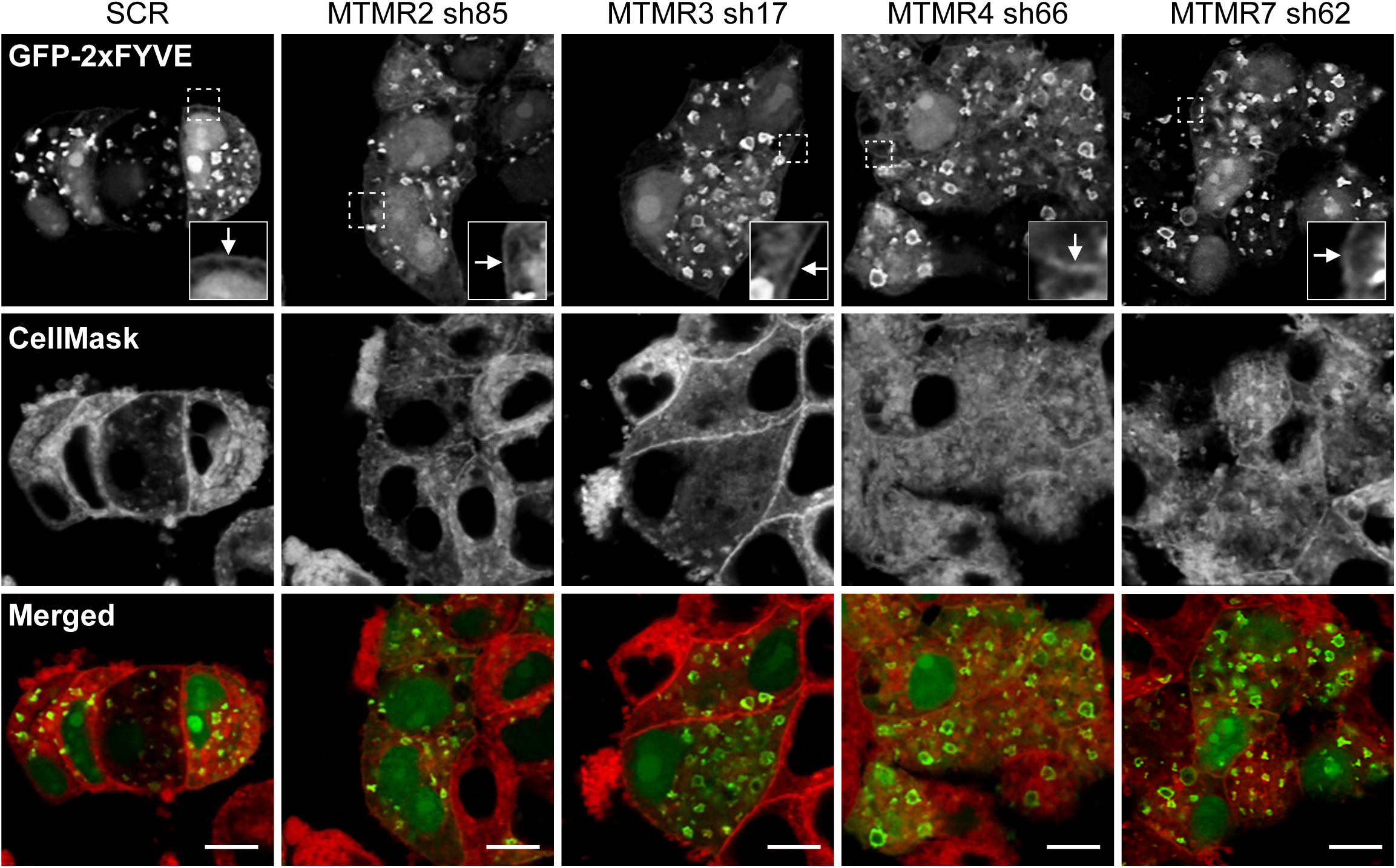
MTMR 2/3/4/7 regulate cellular PI3P contents. T47D cells expressing scrambled shRNA (SCR) or shRNA targeting *MTMR2, 3, 4* or *7*, followed by 1 ug/mL puromycin selection, were overexpressed with GFP-2xFYVE. Cells were incubated with CellMask for 1 h at 37°C incubator, fixed with 4% PFA and imaged by confocal microscopy. A selected region (the white square) is shown at a higher magnification. Arrowheads indicate the PM-staining of GFP- 2xFYVE. Scale bar – 10 μm.

**Supplement Figure 4.**
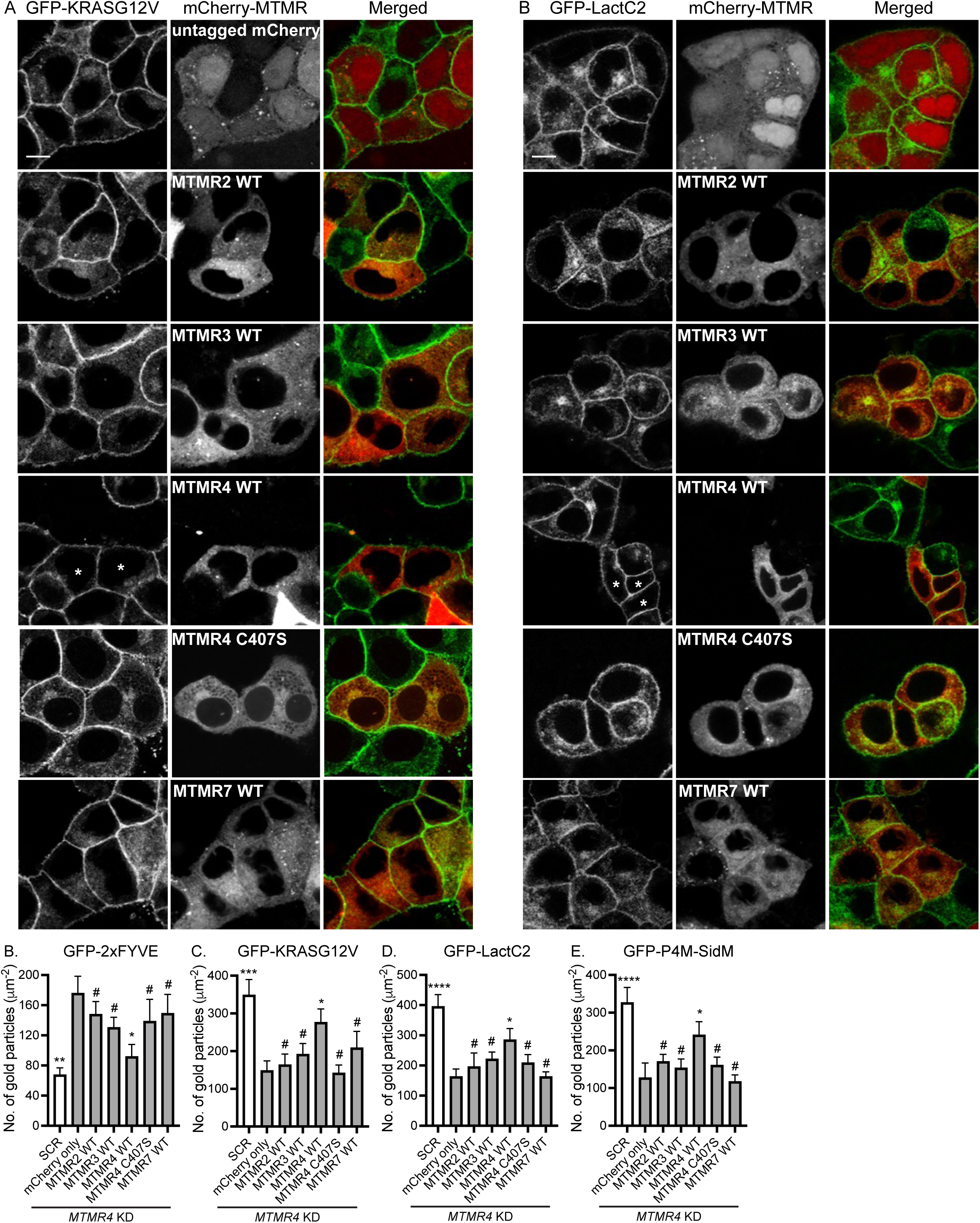
Ectopic expression of WT MTMR4 restores the effects of *MTMR4* loss. T47D cells co-expressing *MTMR4* shRNA66 and GFP-KRASG12V (**A**) or -LactC2 (**B**) were infected with lentivirus expressing indicated mCherry-MTMR proteins, fixed with 4% PFA and imaged by confocal microscopy. Scale bar – 10 μm. Asterisks indicate the restored PM localization of GFP-KRASG12V or -LactC2. Intact basal PM sheets were prepared from T47D cells co-expressing *MTMR4* shRNA66 and GFP-KRASG12V (**C**) or -LactC2 (**D**), -P4M-SidM (**E**), or -2xFYVE (**F**) were infected with lentivirus expressing indicated mCherry-MTMR members, and labeled with anti-GFP-conjugated gold particles and visualized by EM. The graphs show a mean number of gold particles ± S.E.M. (n = 10). Significant differences between control (*MTMR4* KD and untagged mCherry-expressing) cells and other cells were assessed by one-way ANOVA tests for (C – F) (* p < 0.05, ** p < 0.01, *** p < 0.001, # - not significant).

